# Cis-regulatory polymorphism at *fiz* ecdysone oxidase contributes to polygenic adaptation to malnutrition in *Drosophila*

**DOI:** 10.1101/2023.08.28.555138

**Authors:** Fanny Cavigliasso, Mikhail Savitskiy, Alexey Koval, Berra Erkosar, Loriane Savary, Hector Gallart-Ayala, Julijana Ivanisevic, Vladimir L. Katanaev, Tadeusz J. Kawecki

## Abstract

We investigate the contribution of a candidate gene, *fiz* (*fezzik*), to complex polygenic adaptation to juvenile malnutrition in *Drosophila melanogaster*. We show that experimental populations adapted during >250 generations of experimental evolution to a nutritionally poor larval diet (Selected populations) evolved several-fold lower *fiz* expression compared to unselected Control populations. This divergence in *fiz* expression is mediated by a cis-regulatory polymorphism. This polymorphism, which was originally present in a sample from a natural population in Switzerland, is distinct from a second cis-regulatory SNP previously identified in non-African *D. melanogaster* populations, implying that two independent cis-regulatory variants promoting high *fiz* expression segregate in non-African populations. Enzymatic analyses of Fiz protein expressed in *E. coli* demonstrate that it has ecdysone oxidase activity acting on both ecdysone and 20-hydroxyecdysone. Four of five *fiz* paralogs annotated to ecdysteroid metabolism also show reduced expression in Selected larvae, suggesting that malnutrition-driven selection favored general downregulation of ecdysone oxidases. Finally, as an independent test of the role of *fiz* in poor diet adaptation, we show that *fiz* knockdown by RNAi results in faster larval growth on the poor diet, but at the cost of greatly reduced survival. These results imply that downregulation of *fiz* in Selected populations was favored because of its role in suppressing growth in response to nutrient shortage. However, *fiz* downregulation is only adaptive in combination with other changes evolved by Selected populations, such as in nutrient acquisition and metabolism, which ensure that the organism can actually sustain the faster growth promoted by *fiz* downregulation.

## Introduction

Full understanding of adaptive evolution requires elucidation of molecular, developmental and physiological links between specific changes in the genome and fitness-determining traits, such as morphology, behavior or life history. The last two decades brought increasingly detailed insights into a growing number of cases of adaptive evolution mediated by single or few genes. Examples include adaptive variation in color of mammals (Hubbard, et al. 2010), repeated evolution of changes in armor plates in sticklebacks (Colosimo, et al. 2005; O’Brown, et al. 2015), sensory basis of host shift in *Drosophila sechellia* (Auer, et al. 2021), and a trade-off between larval growth and adult flight performance in a butterfly mediated by polymorphism in a respiratory enzyme (Marden, et al. 2013). It is more challenging to identify how specific genetic changes contribute to complex adaptations that involve changes in multiple molecular pathways and physiological mechanisms, and that are mediated by polymorphisms at a large number of genes. Genomics screens and whole genome expression studies have identified numerous candidate polymorphisms and genes involved in traits such as heat tolerance, starvation resistance, fecundity or longevity (Hardy, et al. 2018; Hoedjes, et al. 2019; Jakšić, et al. 2020). However, rather few of these candidates have been verified experimentally (e.g., Martins, et al. 2014; Jakšić, et al. 2020; Hoedjes, et al. 2023), and for most we only have rudimentary understanding of the mechanisms linking genetic variation to phenotypic traits that directly determine fitness.

In this study we explore the contribution of a *cis*-regulatory polymorphism at a single candidate gene to highly polygenic and multifaceted adaptation to juvenile malnutrition in *Drosophila melanogaster*. Nutrient shortage is a fact of life of many animal species, and juveniles in particular are vulnerable, because they usually cannot interrupt their growth and development to wait out for better times. Thus, natural selection imposed by period of nutrient shortage is likely to have shaped animal physiology, with potential consequences for human health (Prentice 2005).

Our study stems from a laboratory evolution experiment in which six *D. melanogaster* populations (subsequently referred to as “Selected” populations) have been raised on a nutritionally poor larval diet for >250 generations (>15 years). In parallel, six Control populations originating from the same base population have been maintained on a standard diet. In the course of this experimental evolution the Selected populations have evolved high tolerance to larval undernutrition, manifested in their higher egg-to-adult survival, faster development and higher larval growth rate on the poor diet compared to Control populations (Kolss, et al. 2009; Erkosar, et al. 2017; Cavigliasso, et al. 2020). This improvement has been in part mediated by an improved efficacy of amino acid acquisition from the poor diet, traded off for reduced rate of absorbing dietary sugars (Cavigliasso, et al. 2020), with associated changes in activity of digestive proteases and amylases (Erkosar, et al. 2017). Metabolome analysis indicated divergence in ways in which the acquired nutrients are used by the Selected and Control larvae, with changes in amino acid and purine metabolism, and an increased use of amino acids for energy generation by Selected larvae (Cavigliasso, et al. 2023). The Selected populations also evolved a smaller critical size at which metamorphosis is initiated, allowing them to complete development with a lower total accumulated biomass (Vijendravarma, et al. 2012a); hence, in spite of their faster larval growth they emerge as smaller adults than Controls (Cavigliasso, et al. 2023). Furthermore, the Selected and Control populations have diverged in their larval foraging behavior (Vijendravarma, et al. 2012b), cannibalistic tendencies (Vijendravarma, et al. 2013) and pre-pupation burrowing (Narasimha, et al. 2015). Consistent with this multitude of phenotypic traits associated with evolutionary adaptation to the poor diet, more than 1300 genes were found to differ in expression between Selected and Control larvae (when both raised on poor diet) (Erkosar, et al. 2017). And, over 3000 single nucleotide polymorphisms (SNPs) in more than 100 genomic regions, in or near about 700 genes, diverged in frequency between Selected and Control populations, implying a highly polygenic genetic architecture (Kawecki, et al. 2021).

Among the candidate genes identified in the RNAseq and SNP scans we focused here on *fiz* (*fezzik*, CG9509, FBgn0030594). With expression several-fold lower in the Selected than Control larvae it was among the genes with the strongest expression divergence (Erkosar, et al. 2017). Furthermore, a non-synonymous SNP in the gene (GLU133LYS), a SNP in the promoter region, and several further SNPs within 3 kb upstream of the gene were significantly differentiated in frequency between the Selected and Control populations in whole genome pool sequencing (Kawecki, et al. 2021). Further inspection of the sequence data revealed that all six Selected populations were fixed or nearly fixed for one allele at each these candidate SNPs, four Selected populations were fixed for the other allele, whereas Control populations C1 and C3 showed intermediate allele frequencies (Figure 1A). The same two Control populations were also intermediate for the level of *fiz* expression (Figure 1B). This concordance between allele frequency and gene expression is suggestive of *cis*-regulatory divergence.

**Figure 1.**
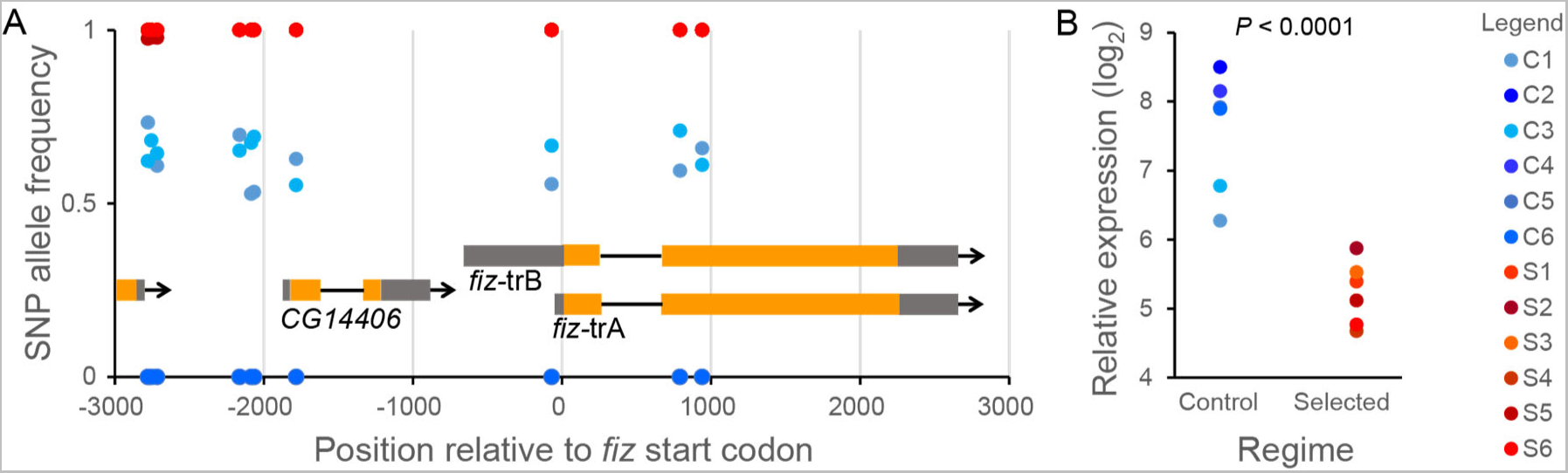
Genomic and expression differentiation of *fiz* gene between Control (C1-C6) and Selected (S1-S6) populations. A. Positions and allele frequencies at candidate SNPs in the genomic vicinity of *fiz* relative to gene transcripts. Allele frequencies are estimated from pooled whole genome sequencing (Kawecki, et al. 2021); for each SNP the frequency plotted is that of the allele more frequent in the Selected than in Control populations. Only SNPs assessed as significantly differentiated between Selected and Control populations are plotted; for a full list of polymorphisms in this region see Supplementary Table S1. B. Expression of *fiz* in third instar larvae raised on the poor diet, based on previously published RNAseq study (Erkosar, et al. 2017). The values are averages from the two conditions used in that study (germ free and *Acetobacter-*inoculated); the expression in the two conditions was highly correlated across populations (*r* = 0.99). *CG14406* is a gene of unknown function immediately upstream; it is not differentially expressed between Selected and Control populations (*P* = 0.75).

*fiz* is mainly expressed in Malpighian tubules (Krause, et al. 2022; UniProt 2023) and, based on homology with *Spodoptera littoralis* ecdysone oxidase and the *Drosophila* gene *Eo*, has been predicted to code for an ecdysone oxidase (Takeuchi, et al. 2005). Ecdysone, or more precisely its active form 20-hydroxyecdysone (20E), is an insect hormone whose pulses trigger transitions between larval instars and the metamorphosis to the adult stage (Kannangara, et al. 2021). However, it also appears to be involved in regulating growth between molts in response to nutrient shortage (Lee, et al. 2018). Ecdysone oxidases catalyze the conversion of ecdysone and 20E to 3- dehydroecdysone (3DE) and 3-epi-20-hydroxyecdysone (3D20E), respectively, the first stage in deactivation of the hormone (Sommé-Martin, et al. 1988; Sun, et al. 2012; Wang, et al. 2018). This makes *fiz* a promising candidate to contribute to adaptive differences between our Selected and Control populations via modulating growth rate and/or critical size and timing of metamorphosis. Another reason why *fiz* is an interesting candidate is a geographic pattern of *cis*-regulatory polymorphism, whereby natural populations in the expanded species range (ranging from North Africa, through Europe to South-East Asia) show 3-4-fold higher expression than ancestral Sub-Saharan populations (Glaser-Schmitt, et al. 2013; Glaser-Schmitt and Parsch 2018). Finally, downregulation of *fiz* has been implicated in microbiota-mediated promotion of larval growth on valine-deficient artificial diet (Grenier, et al. 2023), further suggesting a role in regulating growth response to nutritional stress.

To understand if and how changes in expression of *fiz* might contribute to adaptation to poor diet, we first verified whether Fiz protein indeed has ecdysone oxidase activity. We then characterized the dynamics of *fiz* expression in Selected and Control populations across development and tested whether the differential expression could be compensated by a divergence in expression of other confirmed or putative ecdysone oxidase genes. We tested whether differences in the expression of *fiz* were *cis*-regulatory and aimed to identify the most likely SNP responsible for this divergence. Finally, to independently verify the role of *fiz* in adaptation to poor diet, we investigated how knockdown of *fiz* affects development, growth and survival of larvae on the poor diet.

## Results

### Fiz is an ecdysone oxidase

As the first step towards understanding the potential role of *fiz* in adaptation to larval undernutrition, we verified the bioinformatic prediction that Fiz protein is an ecdysone oxidase, i.e., that it catalyzes the reaction ecdysone + O_2_ *⇌* 3-dehydroecdysone + H_2_O_2_ (Takeuchi, et al. 2005). To verify this prediction, we obtained purified Fiz protein by expressing it in *E. coli and* tested it for oxidase activity using both ecdysone (α-ecdysone) and its activated form 20-hydroxyecdysone (20E, β-ecdysone) as substrates. Incubation of Fiz with either ecdysone or 20E resulted in production of H_2_O_2_ as predicted from the reaction (Figure 2A,B, Supplementary Table S2). Furthermore, in an independent experiment the amount of ecdysone and 20E was reduced when incubated with purified Fiz protein compared to the condition without Fiz (Figure 2C,D, F_1,6_ = 9.7, *P* = 0.021 for ecdysone; F_1,6_ = 9.1, *P* = 0.024 for 20E). Taken together, these results demonstrate that Fiz has an ecdysone oxidase activity and is able to use both ecdysone and 20E as substrates.

**Figure 2.**
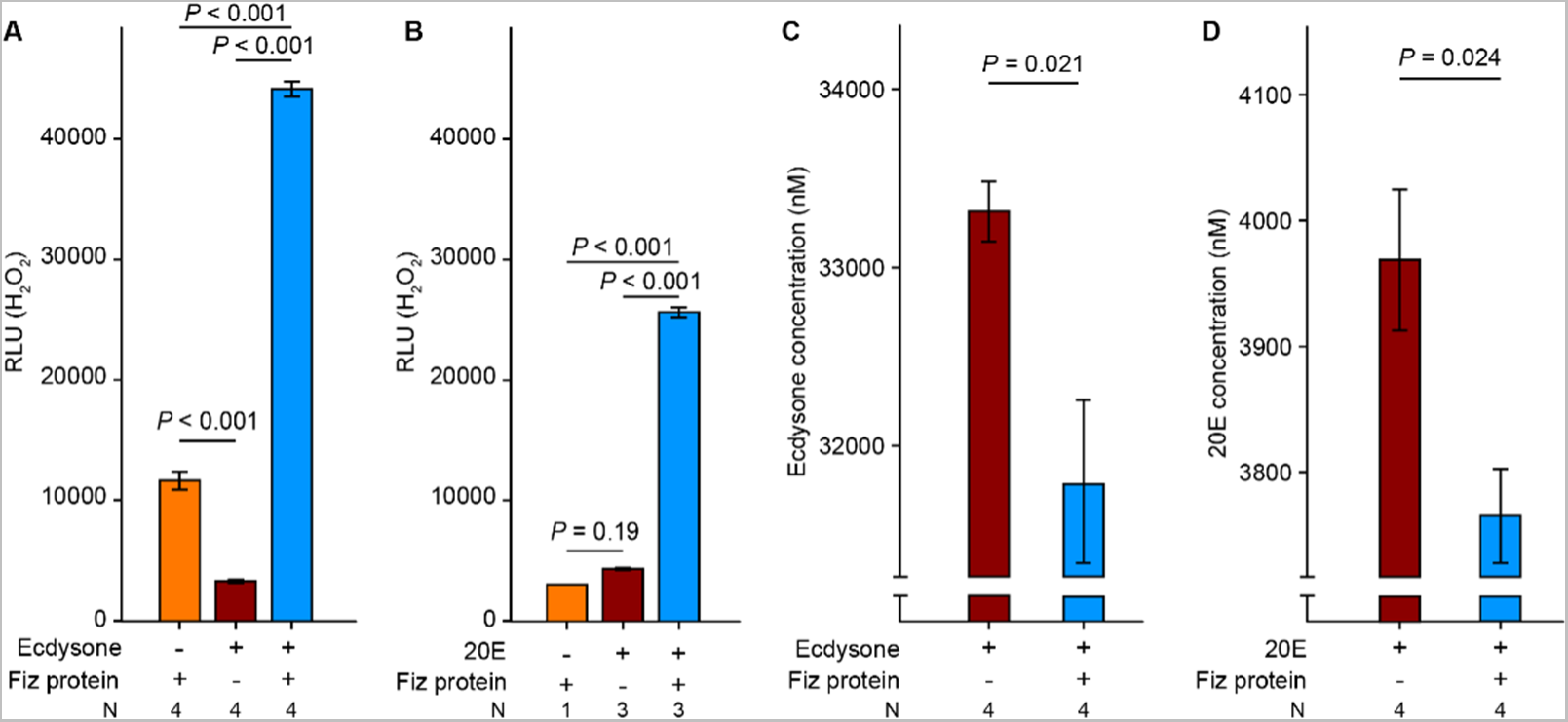
In vitro activity of purified Fiz protein on ecdysone and 20E. A and B. Production of H_2_O_2_ by Fiz protein incubated with ecdysone (A) and 20E (B) as substrates, compared to Fiz protein only or substrate only, in relative units. C and D. Reduction of the relative concentration of ecdysone (C) and 20E (D) following incubation with Fiz protein. Bars correspond to the means ± SE of N = number of replicate reactions; “+” and “-” indicate the presence *versus* absence of a particular compound.

### fiz and its paralogs are less expressed in Selected than Control populations

The RNAseq experiment that detected the difference in *fiz* expression between Selected and Control populations (Erkosar, et al. 2017) was based on a single timepoint during the 3rd larval instar (L3). If *fiz* expression occurred in pulses in response to pulses of ecdysone (Lavrynenko, et al. 2015; Kannangara, et al. 2021), that difference might be due to time shift of the pulse rather than a consistent difference in the level of expression. To address this possibility, we measured the expression of *fiz* at several time points in larvae, prepupae and young adults. To keep the workload manageable, we carried this and the following assays on three Selected (S1, S2, S3) and three Control (C2, C4, C6) populations fixed for alternative alleles at *fiz* candidate SNPs (Figure 1). The expression of *fiz* was consistently lower in these Selected populations compared to Controls, regardless of the developmental stage or diet (Supplementary Figure S1; Supplementary Table S3). In addition, the expression of *fiz* was stable over time in Controls whereas a temporary drop was observed in prepupae of Selected populations.

Of more than a dozen *fiz* paralogs the two with the highest homology to *fiz* (and to *Spodoptera littoralis* ecdysone oxidase) are *Eo*; (FBgn0030597), and *CG9512* (FBgn0030593) (Takeuchi, et al. 2005). Like *fiz* and unlike other paralogs, these two genes are mainly expressed in Malpighian tubules (FlyAtlas2, (Krause, et al. 2022)); the product of *Eo* has been shown to have ecdysone oxidase activity while it had been predicted for *CG9512* (Takeuchi, et al. 2005). The downregulation of *fiz* in Selected populations might thus potentially be compensated by upregulation of these two other ecdysone oxidase genes. This was clearly not the case for prepupae and adults – the expression levels of *Eo* of *CG9512* in Selected populations at these stages were similar to or lower than in Controls (Supplementary Figure S1, Supplementary Table S3). However, the results for the larvae were inconsistent across the timepoints. This could reflect the fact that, because of differences in their rate of development (Kolss, et al. 2009; Erkosar, et al. 2017; Cavigliasso, et al. 2020), the Selected and Control larvae of the same age were at different developmental stages.

We thus quantified the expression of (verified or predicted) ecdysone oxidase genes in larvae that were developmentally synchronized (see Methods), focusing on the first three quarters of the third larval stage between the molt to L3 and the onset of the wandering stage. This is when the pulse of ecdysone that initiates metamorphosis occurs (Koyama and Mirth 2018; Kannangara, et al. 2021). We did this only on larvae raised on standard diet because patterns of expression of ecdysone oxidase genes were broadly similar between diets (Supplementary Figure S1) and because on the poor diet the Selected and Control larvae develop at different rates, and even individuals in the same bottle become desynchronized in their developmental stage. Even though *fiz*, *Eo*, *CG9512* each showed different temporal dynamics, all were consistently less expressed in Selected larvae compared to Controls (Figure 3, *P* < 0.001 for the four genes, Supplementary Table S4). The same was the case for two other *fiz* paralogs with a putative function in steroid metabolism expressed in larvae, *CG9521* and *CG12539* (Figure 3, Supplementary Table S4). A fifth paralog, *CG45065*, a putative ecdysone oxidase mainly expressed in the tracheal system, also tended to be downregulated in Selected mid/late third instar larvae compared to Controls, but this difference was not statistically supported (Figure 3, Supplementary Table S4). Taken together, these results suggest that at least in the first three quarters of the 3^rd^ larval stage, there was no compensation for the downregulation of *fiz* in Selected larvae; rather all known or predicted ecdysone oxidases were (or tended to be) downregulated in the malnutrition-adapted Selected larvae compared to Controls.

**Figure 3.**
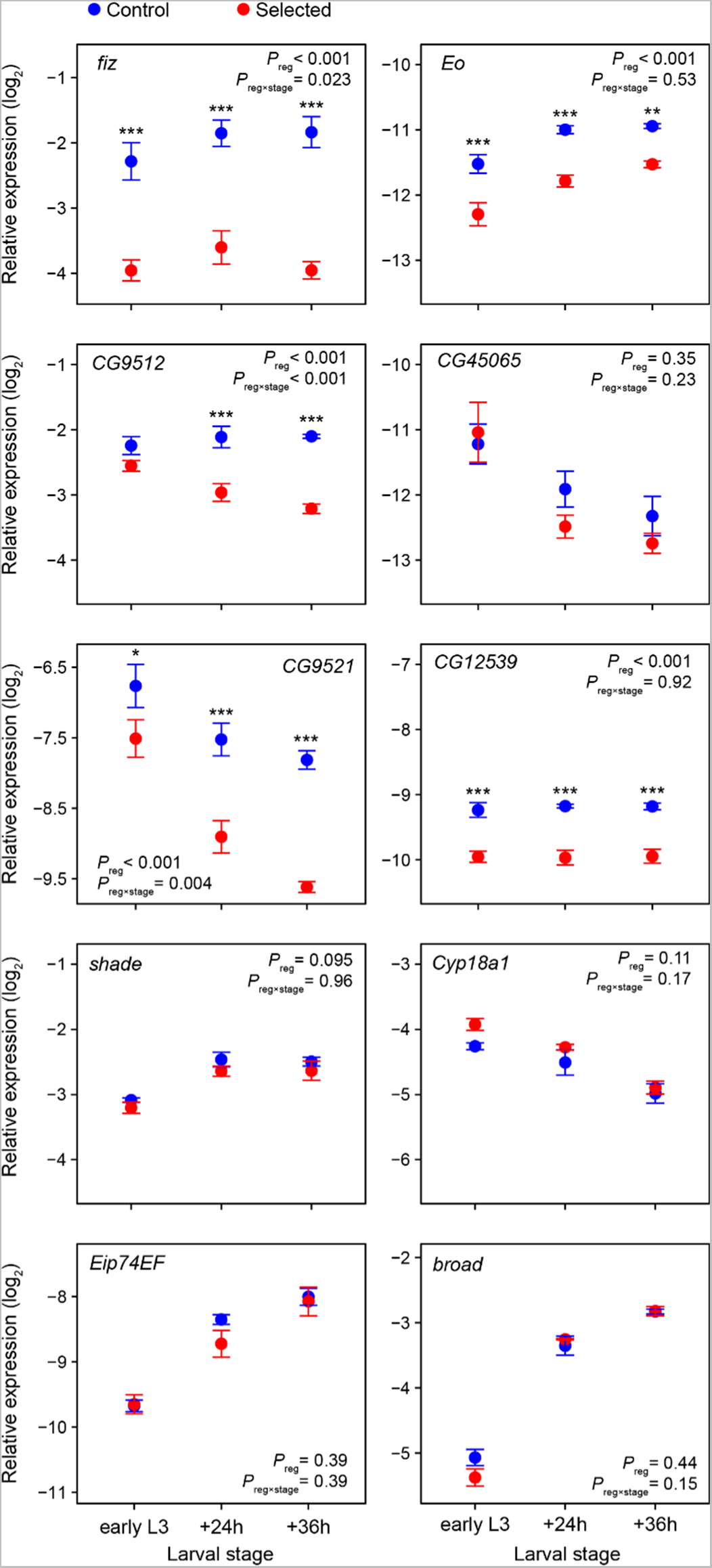
Relative expression of focal genes in third instar larvae raised on standard diet (means ± SE from qPCR relative to three reference genes). Larvae were collected within 17 hours of molting into L3 (“early L3”), as well as 24h and 36h later. For each gene and diet, the *P*_reg_ and *P*_reg×stage_ refer to the main effect of the evolutionary regime (i.e., Selected versus Control) and to regime × stage interaction, respectively. Asterisks indicate a significant stage-specific pairwise difference between Control and Selected populations after *P*-value correction (sequential Bonferroni adjustment for stages). *N* = 3 populations × 3 replicates of 10 pooled larvae per evolutionary regime and stage.

In addition to ecdysone oxidases, products of two other genes are known to directly regulate the active form of ecdysone in peripheral tissues in *Drosophila*: *shade* (*shd*), which converts ecdysone to 20E (Petryk, et al. 2003) and *Cyp18a1* which deactivates 20E by hydroxylation (Rewitz, et al. 2010; Guittard, et al. 2011). We did not detect differences in expression of these two genes between Selected and Control third instar larvae; while the trends observed in the data do not allow us to exclude some differences, they would be much smaller than differences in the expression of ecdysone oxidase genes (Figure 3, Supplementary Table S4).

Based on the putative role of ecdysone oxidases in deactivating ecdysone, one would thus expect that their lower expression in Selected larvae would lead to a higher level of 20E. The concentration of 20E in third instar *Drosophila* larvae is below or at detection threshold until shortly before pupariation (Lavrynenko, et al. 2015), long after the (relatively low) peak of 20E that initiates metamorphosis (Kannangara, et al. 2021). The level of 20E at this stage is thus often quantified indirectly by measuring expression of genes activated by 20E, such as *Eip74EF* and *broad* (Karim and Thummel 1991; Glaser-Schmitt and Parsch 2018; Texada, et al. 2019). Expression of both *Eip74EF* and *broad* increased in the course of the third larval instar, consistent with the expected increase in 20E levels. However, the expression of these two genes did not differ between Selected and Control larvae and, if anything, tended to be lower in Selected larvae at individual time points (Figure 3, Supplementary Table S4). Thus, we have no indication of 20E reaching higher levels in Selected larvae, despite lower expression of *fiz* and other ecdysone oxidase genes. This result is inconsistent with degradation of ecdysone and 20E being the sole function of ecdysone oxidases in *Drosophila*, a point we elaborate on in Discussion.

### Difference in fiz expression is cis-regulatory and not sex-specific

The existence of candidate SNPs upstream from *fiz* opens the possibility that the difference in expression between the Selected and Control populations is mediated by one (or more) of these variants in a *cis*-regulatory manner. If so, individuals (females because *fiz* is on the X) carrying one chromosome each from a Selected and a Control population would express the *fiz* copy on the “Selected” chromosome much less the copy on the “Control” chromosome. Based on pool sequencing data (Kawecki, et al. 2021), and verified at the time of the present experiment, the Control populations C2, C4 and C6 and the Selected populations S1, S2, S3 are fixed for alternative alleles at a SNP in the coding region of *fiz* (G/A at position 791 after start codon). This allowed us to quantify the relative contribution of the two alleles to the pool of *fiz* transcripts in heterozygotes with cDNA amplicon sequencing. We did this in F1 female late 3rd instar larvae from three independent crosses (in both directions) between Control and Selected populations (C2 × S1, C4 × S2 and C6 × S3). Regardless of the direction of the cross, these F1 females showed a highly allele-biased expression, with the transcript of the “Control” allele “A” being over 5-fold more abundant than the transcript of the “Selected” allele “G” (Figure 4A, t_4_ = 31.8, *P* < 0.001, Supplementary Table S6). This asymmetry in expression was nearly as large as in samples consisting of a 50:50 mix of Selected and Control cDNA (“Mix” in Figure 4A). This implies that most – although possibly not all – of the difference in *fiz* expression between Selected and Control populations is *cis*-regulatory.

**Figure 4.**
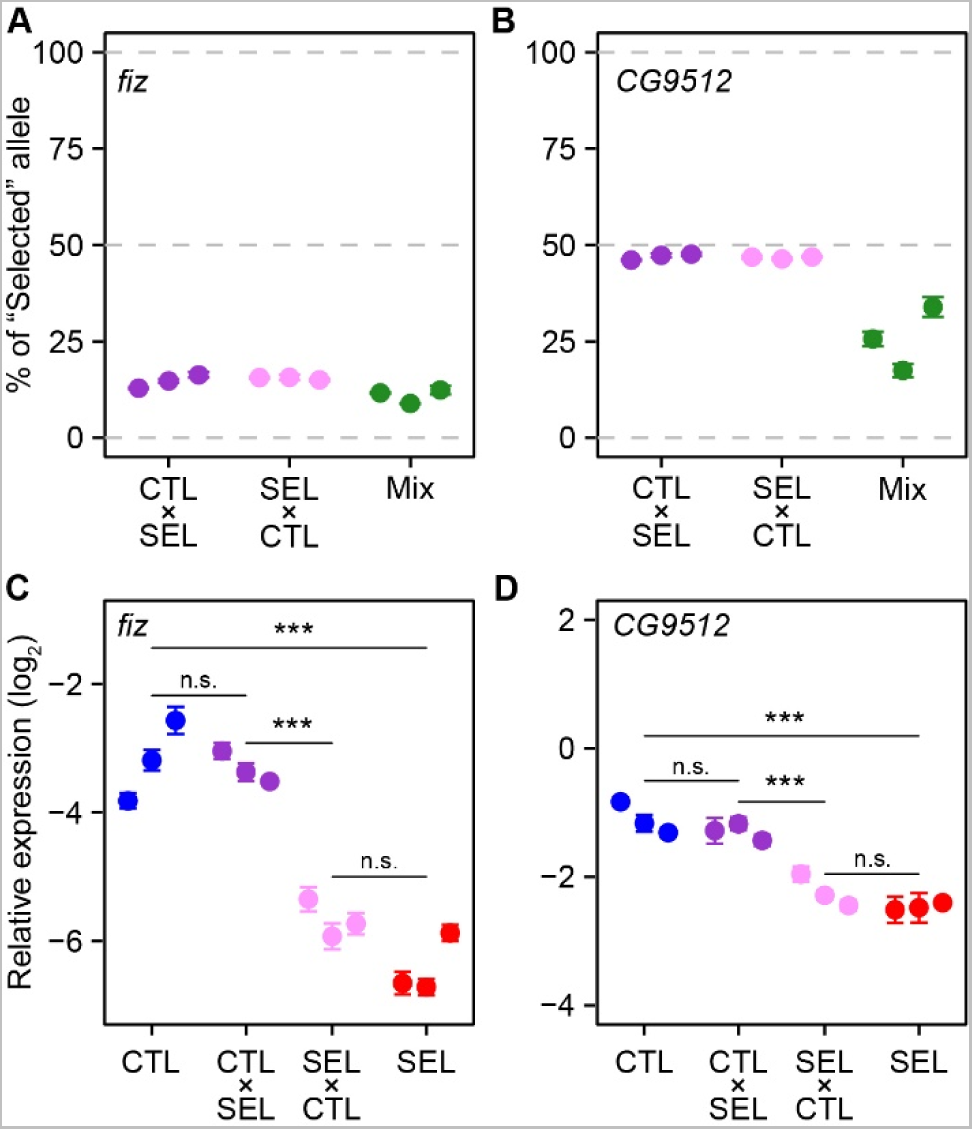
Expression of alleles of *fiz* and *CG9512* in F1 crosses between Control and Selected populations. A,B: Proportional contribution of the “Selected” allele to the transcript pool of *fiz* (A) and *CG9512* (B) in heterozygous F1 female larvae, determined by amplicon sequencing. The proportion of the “Selected” allele in 50:50 mix of cDNA from parental lines provides a reference. C,D: Relative expression (from RT-qPCR) of *fiz* and *CG9512* in F1 male larvae (hemizygous for both X chromosome genes) compared to the parental populations. SEL: Selected; CTL: Control; F1 crosses represented as maternal × paternal population. The three symbols per category correspond to three independent population pairs (C2 × S1, C4 × S2, C6 × S3). Each symbol indicates mean ± SE of N = 2 to 4 (A-B) or 4 (C-D) replicates per population or cross; each replicate consisted of a pool of seven late third instar larvae. \*\*\**P* < 0.001, n.s. *P* > 0.05.

We used the same approach to test for *cis*-regulatory nature of differences in gene expression in another (predicted) ecdysone oxidase, *CG9512*. Its transcript begins less than 1kb downstream from the end of *fiz*, it is highly expressed in larvae (Figure 3) and it also harbors a SNP (T/C) in the coding region of *fiz* (position 904 after start codon) fixed for alternative alleles in the three Selected from the three Control populations. Even though the Selected allele tended to be slightly less abundant than the Control allele in the *CG9512* transcript pool, the difference was much smaller than that between their relative abundance in the 50:50 mix of Selected and Control larvae (Figure 4B, Supplementary Table S6). Thus, in contrast to *fiz*, the difference in expression of *CG9512* appears mainly *trans*-regulatory.

As *fiz* and *CG9512* are on the X-chromosome, males only inherit the maternal copies of these genes. For *fiz*, consistent with *cis*-regulatory nature of the expression difference, male larvae of crosses between Selected and Control populations showed *fiz* expression similar to that of their maternal population and very different from each other (Figure 4C, Supplementary Table S7). These results also confirmed that larvae of both sexes showed a similar difference in *fiz* expression, validating results obtained without sexing the larvae (which is impractical before mid-3rd larval stage). As for *fiz*, the expression of *CG9512* also depended on the direction of the cross, with the expression in each cross being similar to that in its maternal population (Figure 4D, Supplementary Table S7). While in principle this could mean that, in contrast to females, the difference in *CG9512* expression in males is *cis*-regulatory, a more parsimonious explanation is that the expression difference is mediated by a *trans*-regulatory element located on the X-chromosome.

### The responsible polymorphism is distinct from previously identified cis-regulatory SNP

We examined previously published pool seq data (Kawecki, et al. 2021) for candidate polymorphisms potentially responsible for these *cis*-regulatory differences in *fiz* expression. A previous study identified a *cis*-regulatory C/G SNP at genomic position X: 14909071 with a large effect on *fiz* expression; Sub-Saharan populations are fixed for the “low-expression” ancestral allele C whereas cosmopolitan populations harbor a derived “high expression” allele G at frequencies ranging from 0.18 to 0.50 (Glaser-Schmitt, et al. 2013; Glaser-Schmitt and Parsch 2018). This polymorphism (referred to as “SNP 67” in (Glaser-Schmitt and Parsch 2018)) is absent in our populations – all are fixed for the “low expression” C allele (Kawecki, et al. 2021).

Hence, the divergence in *fiz* expression in our study must be mediated by an alternative *cis*- regulatory polymorphism. A likely alternative candidate is a G/T SNP just 5 bp away, at genomic position X:14909076, 70 bp upstream of the start codon and within a predicted *fiz* promoter (Eukaryotic Promoter Database, Dreos, et al. (2015); Meylan, et al. (2020)). At this SNP all Selected populations are fixed for the G allele, Control population C2, C4, C5, C6 are fixed for the T allele, and populations C1 and C3 – which show intermediate *fiz* expression - harbor intermediate frequencies (Figure 1, Supplementary Table S1). This results in a nearly perfect correlation across populations between allele frequency at this SNP and *fiz* expression (*r* = 0.96, *P* < 0.001). This polymorphism has been reported before, with the G allele being fixed in Sub-Saharan populations and presumably ancestral, and the derived T allele being found at low to moderate frequencies in many populations across Europe, North Africa and Asia (Glaser-Schmitt, et al. 2013; Kapun, et al. 2020).

Ten other SNPs and three short (2-7 bp) indels had been found within 1 kb upstream of the *fiz* gene (and further seven SNPs in the intron of the gene, Supplementary Table S1), but none of them was identified as a candidate for targets of selection by Kawecki, et al. (2021). However, *fiz* lies within an apparent 355 kb genomic block containing 218 candidate SNPs, most of which show similar patterns of allele frequencies to the SNP in *fiz* promoter (Figure 2C and Supplementary Table S3 in Kawecki, et al. (2021)). Eleven such candidate SNPs are located between 1.7 and 3.5 kb upstream of *fiz* (Figure 1A). Thus, even though the SNP in *fiz* promoter is a highly promising candidate, we cannot exclude that the *cis*-regulatory element responsible for differences in *fiz* expression in our populations is farther away from the gene. However, these differences in expression are clearly not mediated by the SNP previously identified by Glaser-Schmitt and Parsch (2018).

### fiz knockdown promotes growth but impairs survival on poor diet

The *cis*-regulatory divergence in *fiz* expression between the Selected and Control populations is suggestive of its contribution to dietary adaptation. However, *fiz* lies within a genomic region where many SNPs show highly parallel changes in allele frequency, presumably due to linkage disequilibrium (Kawecki, et al. 2021). This region contains about 40 genes. Hence, rather than being favored by selection, the divergence between Selected and Control populations in the expression of *fiz* may have evolved as a byproduct on selection on linked polymorphisms affecting expression or functioning of other genes. Therefore, as an independent test of the role of reduced *fiz* expression in adaptation to undernutrition, we studied the effect of RNAi knockdown of *fiz* on larval performance traits. *fiz* knockdown has been previously shown to promote larval growth on a nutrient-rich diet (Glaser-Schmitt and Parsch 2018). If *fiz* expression divergence between the Selected and Control populations were adaptive, we would expect *fiz* knockdown to have positive effects on performance on at least some aspects of performance on the poor diet.

The knockdown considerably reduced egg to adult survival on both poor and standard diet (Figure 5A; χ^2^ = 17.7, *P* < 0.001 and χ^2^ = 14.3, *P* < 0.001, respectively). It aslo accelerated egg-to-adult development of both sexes on poor diet (Figure 5B, F_1,6_ = 18.9, *P* = 0.005, Supplementary Table S8), but not on standard diet (Figure 5B, F_1,8_ = 0.3, *P* = 0.59, Supplementary Table S8). Furthermore, regardless the diet, *fiz* knockdown flies were larger in terms of body mass, and this was not mediated by prolonged development; rather, for any length of development, the knockdown females were heavier than the controls (Figure 5C; F_1,11_ = 25.1, *P* < 0.001 on poor diet; F_1,21_ = 13.0, *P* = 0.002 on standard diet, Supplementary Table S9). This implies that the *fiz* knockdown improved the growth rate: even though (as is typical for *Drosophila*) slower development was associated with slower growth, for any length of development the estimated growth rate of knockdown larvae was higher than that of controls (Figure 5D, Supplementary Table S10).

**Figure 5.**
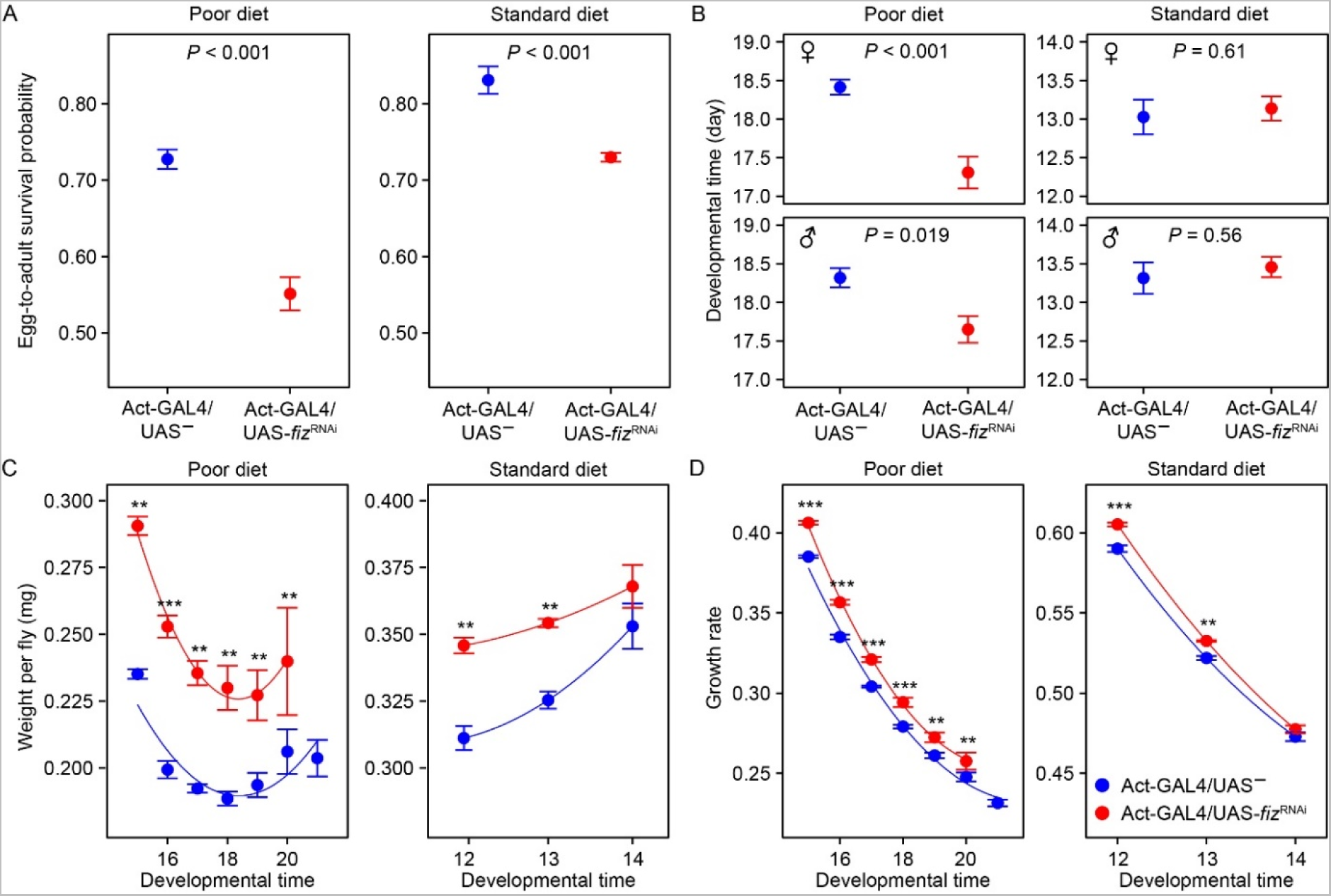
Effect of *fiz*-knockdown on egg-to-adult survival probability (A), developmental time (B), female weight (C) and female growth rate (D) on poor and standard diets. Symbols indicate means ± SE. *N* = 4 replicate bottles for poor and 5 bottles for standard diet (each bottle initially contained 200 eggs). Red and blue lines in C and D represent predictions from statistical models (including the linear and quadratic components).

This experiment thus demonstrated that a reduction of *fiz* expression by RNAi in a non-adapted genetic background was sufficient to improve larval growth rate and thus adult size on both diets. The effects were greater on the poor than standard diet (Supplementary Table S11), and development was accelerated only on poor diet. However, this happened at the expense of reduced survival on both diets, possibly explaining why a low expression of *fiz* had not been favored in Control populations.

## Discussion

### Enhanced fiz expression evolved twice in out-of-Africa populations

We discovered a several-fold divergence in the expression of *fiz* between populations of *D. melanogaster*, driven by laboratory natural selection imposed by diet quality and mediated by a *cis*- regulatory polymorphism. Previous studies reported a geographical pattern of variation in *fiz* expression in *D. melanogaster*. While populations from the ancestral range in sub-Saharan Africa show low expression, many cosmopolitan populations (from Europe, North Africa, Central and South-East Asia) contain a substantial frequency of genetic variants with several-fold higher *fiz* expression (Hutter, et al. 2008; Glaser-Schmitt, et al. 2013; Glaser-Schmitt and Parsch 2018). Increased *fiz* expression is associated with reduced adult size and reduced wing loading, and this may have facilitated *D. melanogaster* expansion out of Africa (Glaser-Schmitt and Parsch 2018). However, most cosmopolitan populations have remained polymorphic, with both high- and low-expression variants likely maintained by a combination of fluctuating and sexually antagonistic selection (Glaser-Schmitt, et al. 2021). In the course of evolutionary adaptation to a nutrient-poor larval diet, our Selected populations reverted to the Sub-Saharan low expression phenotype of *fiz*, while the Control populations maintained on standard diet retained the polymorphism or became fixed for the high expression variant.

A single nucleotide polymorphism in the promoter region has previously been identified as responsible for much of variation in *fiz* expression (genome reference position X:14909071, or “SNP 67”; Glaser-Schmitt and Parsch 2018). However, all our populations are fixed for the low expression variant at this position. The divergence in *fiz* expression between Selected and Control populations must therefore be mediated by a different genetic polymorphism, a likely candidate being the SNP within *fiz* promoter at position X:14909076. Thus, irrespective of the forced favoring it, increased *fiz* expression appears to have evolved independently twice after out-of-Africa expansions of *D. melanogaster*, a case of convergent evolution at the molecular level. Such cases of independent evolution of the same molecular phenotype based on different mutations of the same gene are rather scarce in animals (e.g., Chan, et al. 2010; Cosme, et al. 2020).

### Downregulation of fiz reverses undernutrition-induced growth inhibition

We found that *fiz* knockdown results in faster larval growth on a nutrient-poor diet. This is likely the reason (or at least a major reason) why downregulation of *fiz* was favored in the course of experimental evolution of our Selected populations on the poor diet. Faster growth under nutritional restriction can be mediated by two complementary mechanisms: enhanced acquisition of dietary nutrients resulting in greater availability of raw materials (amino acids, lipids, etc.) and energy, and promotion of growth via signaling pathways that regulate the rate at which these raw materials and energy are used for growth. While adaptation of the Selected populations to undernutrition involves both mechanisms (Erkosar, et al. 2017; Cavigliasso, et al. 2020; Cavigliasso, et al. 2023), our results and preexisting knowledge indicate that *fiz* downregulation contributed to their adaptation via the latter mechanism, i.e., changes in growth regulation. *fiz* has been identified as growth inhibitor in *Drosophila* larvae (Glaser-Schmitt and Parsch 2018); in particular it mediates growth inhibition by diet with unbalanced amino acid content (Grenier, et al. 2023). We confirmed the bioinformatic prediction that *fiz* encodes an oxidase of both ecdysone and 20E. This suggests that *fiz* acts via modulating ecdysone signaling. Ecdysone is a key insect hormone that triggers molts and metamorphosis; however, it also acts as a negative regulator of systemic growth throughout development, in particular in response to nutrient shortage (Colombani, et al. 2005; Delanoue, et al. 2010; Sun, et al. 2014; Kannangara, et al. 2021) and under mild hypoxia (Kapali, et al. 2022). It is unlikely that the downregulation of *fiz* contributes to better nutrient acquisition. Ecdysone or ecdysteroids in general are not known to mediate nutrient acquisition, and the high mortality of *fiz* knockdown larvae on the poor diet suggests that their fast growth drains nutrients and energy from life sustaining processes. In other words, *fiz* knockdown on its own appears to perturb an adaptive adjustment of growth rate to nutrient availability.

### Ecdysone oxidases may do more than deactivate ecdysone

We demonstrated that, consistent with bioinformatic prediction, purified Fiz protein has ecdysone oxidase activity in vitro, acting on both ecdysone and 20E. We found that, in addition to *fiz*, four out of five *fiz* paralogs expressed in larvae annotated as oxidoreductases involved in ecdysteroid metabolism were also downregulated in Selected larvae compared to Controls (Figure 2). This suggests that selection imposed by the poor diet generally acted to reduce ecdysone oxidase activity. However, our results are not consistent with deactivation of ecdysone and 20E being the sole function of *fiz* and, by extension, of other ecdysone oxidases in *Drosophila*, an inference corroborated by several published studies. The notion that oxidation of ecdysone and 20E is the first step in their degradation mainly comes from research on Lepidoptera, in which a further reaction leads to epimerization of these ecdysteroids, followed by excretion (Takeuchi, et al. 2001; Sun, et al. 2012; Rees 2013). Consistent with this, high expression of ecdysone oxidase in silkworm is associated with low levels of 20E and vice versa (Sun, et al. 2017; Wang, et al. 2023). If this result could be extrapolated to *Drosophila*, reduced ecdysone oxidase activity in Selected populations should lead to higher levels 20E. This does not appear to be the case, at least in the early to mid-third instar larvae– we did not detect any differences in the expression of genes that respond to the ecdysone peak that triggers the processes leading to metamorphosis (*Eip74EF* and *broad*). In another study, a null mutant of *fiz* was reported to have a lower expression of *Eip74EF* and a couple of other 20E-induced genes in late 3rd instar (Glaser-Schmitt and Parsch 2018), suggesting that *fiz* knockout actually leads to a lower 20E titer, contrary to what would be predicted if *fiz* acted to deactivate 20E. The notion that *fiz* deactivates 20E also does not mesh up the fact that 20E acts as growth inhibitor (Colombani, et al. 2005; Delanoue, et al. 2010; Kapali, et al. 2022). If the main function of *fiz* were to deactivate 20E, *fiz* knockdown should thus result in slower growth, whereas the opposite is the case.

These considerations raise the question of the function(s) of ecdysone oxidases. First, there seems to be little evidence that epimerization is a major route of ecdysteroid deactivation in *Drosophila* (Rewitz, et al. 2013), and no gene has been annotated to the enzyme performing the second step in this pathway (3DE-3α-reductase) (https://www.genome.jp/pathway/dme00981). Instead, 20E deactivation, at least during metamorphosis, appears to occur chiefly via hydroxylation catalyzed by Cyp18a1 (Rewitz, et al. 2010; Guittard, et al. 2011). Second, several lines of published evidence suggest that products of ecdysone oxidase acting on ecdysone and 20E (3DE and 3D20E, respectively) are more than deactivated compounds on the way to degradation or excretion. In particular, 3D20E activates the ecdysone receptor (EcR), albeit to a lesser degree than 20E (Baker, et al. 2000). Sommé-Martin, et al. (1990) reported that 3D20E is more potent than 20E to induce the expression of an ecdysone-dependent gene *Fbp1* in *Drosophila* larvae. In silkworm, 3DE produced in ovaries appears to serve as an inactive reservoir of ecdysteroids that is converted rapidly to 20E during embryonic development (Wang, et al. 2018). Such “recycling” of ecdysone and 20E deactivation products has also been proposed as a major source of 20E peak during a late stage of *Drosophila* metamorphosis, which occurs after the larval prothoracic gland has disintegrated (Scanlan, et al. 2023). Our results suggest the intriguing possibility that the products of ecdysone/20E oxidation directly play an active role in regulating larval growth in *Drosophila* or can be recycled into active compounds that do so.

### Adaptation to malnutrition involves coadapted traits

The adaptation of Selected populations to poor diet is manifested in both faster larval growth and higher egg-to-adult survival compared to Controls (Kolss, et al. 2009; Erkosar, et al. 2017; Cavigliasso, et al. 2023). In contrast, even though *fiz* knockdown resulted in substantially enhanced larval growth on the poor diet (thus recapitulating one aspect of adaptation in Selected populations), this occurred at a great cost to survival. This suggests that a reduction in *fiz* expression on its own is maladaptive and improves fitness on the poor larval diet only in combination with evolutionary changes in other traits, mediated by other genes. As discussed above, our and other authors’ results indicate that *fiz* inhibits growth. If so, its downregulation in the Selected larvae would allow them to better realize their growth potential despite nutrient shortage, potential that has been enhanced owing to better assimilation of dietary amino acids (Cavigliasso, et al. 2020) and changes in amino acid metabolism (Cavigliasso, et al. 2023). In contrast, downregulation of *fiz* in otherwise non-adapted larvae appears to cause them to overextend themselves trying to grow at a rate that their metabolism cannot sustain. Furthermore, the faster growth of *fiz*-knockdown larvae compels them to accumulate more biomass before attempting metamorphosis, which becomes increasingly challenging as the condition of the diet deteriorates over time as the nutrients are extracted and metabolic waste products accumulate. Such attempts to grow beyond what the available means permit might explain their high mortality. In contrast, owing to their reduced critical size for metamorphosis initiation, the Selected larvae convert their faster growth into fast development and actually emerge smaller as adults than Controls (Vijendravarma, et al. 2012a; Cavigliasso, et al. 2023). Thus, only with these other facets of adaptation to undernutrition is a reversal of *fiz*-mediated growth inhibition adaptive. Thus, while the precise mechanisms remain to be elucidated, our results indicate that adaptation to larval undernutrition is not only polygenic, but also involves epistatic interactions and multiple coadapted traits.

## Materials and Methods

### Synthesis and purification of Fiz protein

We amplified the coding region of the *fiz* gene from cDNA clone (FI24119, *Drosophila Genomics Resource Center)* with primers (forward:

GATCTCACCATCACCATCACCATTCACTAGACGGTGGCCAGAAT; reverse:

AAGCTTGTCGACGGAGCTCGGCCAATCGAAACGAGTTCTGTGT) using Phusion High-Fidelity DNA Polymerase (New England Biolabs, Ipswich USA, cat. #M0530S). The vector pET-23b_RGS_6xHis_Go (Lin, et al. 2014) was used for subcloning amplified *fiz* sequence, and this vector was digested with NcoI and EcoRI to remove Go sequence. We purified the vector by gel extraction using GeneJet Gel extraction kit (K0692, ThermoFisher Scientific™, USA). NcoI-EcoRI fragment containing backbone plasmid and N-terminal 6xHis tag was assembled with amplified *fiz* sequence. Cloning step was performed using the NEBuilder HiFi DNA Assembly Cloning Kit (New England Biolabs, cat. #E5520S). The cloning was verified by restriction analysis and sequencing.

We transformed *Escherichia coli* strain DE3-Rosetta Gami with our plasmid (pET-23b_RGS_6xHis_fiz) and grew overnight at 37°C on a Petri dish. Pre-culture initiated with two independent colonies were then grown overnight at 37°C in LB solution. These precultures were mixed 1:1, and we subsequently inoculated a small volume of this pre-culture with a mix of 1 × LB, 100 μg/mL ampicillin, 10 μg/L polypropylene glycol (pre-culture:medium ratio of 1:100) and grew at 37°C and 180 rpm to reach an OD_600_ = 0.6-0.8. Cultures were then cooled down for 30 min at 17°C before a further 8h overnight incubation at 17°C with 1 mM isopropyl-1-thio-d-galactopyranoside (IPTG). After 10 min centrifugation at 6000 × g, we resuspended cell pellets in 1 × TBS (20mM Tris-HCl pH 7.4, 150mM NaCl) and 1 mM PMSF, before lysing them with a Cell disruptor (Constant systems). We removed debris by centrifugation at 15,000 × g/15 min/4°C. We collected and mixed the supernatant with 30 mM imidazol and 50 μl of drained beads (HisPur Ni-NTa Resin) per liter of initial culture before incubating at RT for 1 h on a rotating shaker and centrifuging at 1,000 × g/1 min/4°C. The pellet was washed five times. Each washing consisted of adding 50 mL of washing buffer (1 × TBS and 10 mM imidazole) before incubation on ice for 10-15 min to settle the beads and remove the supernatant. For the third washing, the 10 min incubation on ice has been replaced by an incubation at 4°C for 1 h on a rotating shaker. The elution was done with 1 × TBS and 300 mM imidazole. The elution volume corresponded to two volumes of drained beads. After a short centrifugation, the supernatant was collected, and the purity of the protein was checked on a gel (Supplementary Figure S2). We then performed a Bradford assay (Roti®-Quant, K015.3, Roth) to estimate the protein concentration. Fiz being predicted to be a flavoprotein (UniprotKB, automatic annotation, (UniProt 2023)), we confirmed that our purified Fiz contains FAD (Supplementary Table S12) which is a proxy for the proper folding of flavoproteins.

### Enzymatic activity of Fiz protein in vitro

Ecdysone oxidases (KEGG EC 1.1.3.16) use O_2_ to convert ecdysone substrate into 3-dehydroecdysone and H_2_O_2_; this reaction is thought to deactivate the hormone (Takeuchi, et al. 2005). No chemical standard of 3-dehydroecdysone was available, precluding an assay based on its detection/quantification. Thus, to test whether Fiz has ecdysone oxidase activity, we used two other methods, one based on quantifying H_2_O_2_ production with a fluorescent enzymatic assay and the other using liquid chromatography – mass spectrometry (LC-MS) to measure the decrease of the ecdysone substrate. In addition to ecdysone (a.k.a. α-ecdysone, KEGG C00477), in both methods we also used the active form of ecdysone, 20-hydroxyecdysone (20E, a.k.a β-ecdysone or β- ecdysterone; KEGG C02633) as a potential substrate.

For the first method, the H_2_O_2_ production was quantified with ROS-Glo^TM^ H_2_O_2_ assay kit (#G8820, Promega, Switzerland). We incubated 28 μg of purified Fiz with 20 μL of 125 μM ecdysone or 20E and Tris-HCl (pH = 8) to reach a final volume of 80 μL, and with 20 μl of H_2_O_2_ substrate from the kit for 1 h. We then incubated the reaction with ROS-Glo detection solution for 20 min. The amount of H_2_O_2_ was estimated by measuring relative luminescent values with a plate reading luminometer (Hidex Sense Plate Reader, Labgene, Switzerland). Two negative controls were done for each assay: one with the substrate (Ecdysone or 20E) and without the Fiz protein (buffer only), and one with the Fiz protein and without the substrate. Four and three replicate reactions were quantified for each condition involving Ecdysone or 20E, respectively. Five negative controls with only Fiz protein (no ecdysone/20E) were performed, four simultaneously with the Ecdysone assay and together with the 20E assay.

For the second method, we incubated 28 μg of purified Fiz with 20 μL of ecdysone 125 μM or 20E 125 μM and Tris-HCl pH 8 (final volume of 80 μL) for 80 min. As negative control, we replaced Fiz by its TBS buffer. The reaction was then quenched with 320 μL of 80% methanol and stored at –80°C. Alpha-ecdysone and 20-hydroxyecdysone (beta-ecdysone) were quantified by LC-MS/MS analysis in positive ionization mode using a TSQ Altis triple quadrupole system (QqQ) interfaced with Vanquish Horizon UHPLC system (Thermo Scientific). The chromatographic separation was carried out in a Accucore^TM^ aQ column (2.6 μm, 100 mm × 2.1 mm I.D) (Thermo Scientific). Mobile phase was composed of A = 0.1 % acetic acid in water and B = 0.1% acetic acid in ACN at a flow rate of 250 μL/min. Column temperature was 40 °C and sample injection volume 2µL. The linear gradient elution starting from 2% to 100% of B (in 5 min) was applied and held until 6 min. The column was then equilibrated to initial conditions. ESI source conditions were set as follows: voltage 3500 V in positive mode, Sheath Gas (Arb) =50, Aux Gas (Arb) =10, Sweep Gas (Arb) =1 and Ion Transfer Tube Temperature 275 °C. Nitrogen was used as the nebulizer and Argon as collision gas (1.5 mTor). Vaporizer Temperature was set to 350 °C. Selected Reaction Monitoring (SRM) was used as acquisition mode with a total cycle time of 400 ms. Optimized collision energies for each metabolite were applied. Raw LC-MS/MS data was processed using the Thermo Sceintific Xcalibur software. External calibration curves (spanning the wide concentration range from 1nM to 20µM) were used to report the estimated concentrations of ecdysone and 20E.

The activity of purified Fiz was analyzed by fitting a linear model (LM) with either the amount of H_2_O_2_ or substrate (Ecdysone or 20E) as response variable and the condition (purified protein + substrate (E or 20E) or purified protein only or substrate only (E or 20E)) as a fixed factor. We used a type 2 F-test and when applicable, pairwise comparisons were performed with *emmeans* and *pairs* functions in R.

### Experimental evolution

This study is based on a long-term laboratory evolution experiment, in which six Selected populations of *Drosophila melanogaster* have been evolving on a nutritionally poor larval diet for over 15 years, with six Control populations maintained in parallel on standard diet. These populations were originally derived from a lab-adapted base population collected in Basel, Switzerland in 1999 and subsequently maintained in the lab on the standard diet (15 g agar, 30 g sucrose, 60 g glucose, 12.5 g dry brewer’s yeast, 50 g cornmeal, 0.5 g CaCl2, 0.5 g MgSO4, 10 mL Nipagin 10%, 6 mL propionic acid, 20 mL ethanol per liter of water) until the evolution experiment commenced in 2005. The Control populations have continued to be maintained on the standard diet. The six Selected populations have been maintained on a diluted larval diet, containing one-fourth of the amounts of sugars, yeast and cornmeal of the standard diet; we refer to this diet as “the poor diet”. All populations were maintained at a controlled density of approximately 200-250 eggs for 40 mL of food, with a generation cycle of 3 weeks and census size of about 180-200 breeding adults. After their emergence, adult flies of both selection regimes were transferred to standard diet and additionally fed ad libitum live yeast to allow mating and egg laying. Fly stock maintenance and all experiments were performed at 25°C with 50–70% humidity and 12:12 light cycle. Details of the fly maintenance are described in Kolss, et al. (2009).

For this study, we chose a subsample of three Control (C2, C4, C6) and three Selected (S1, S2, S3) populations. This allowed us to limit the workload so that despite multiple timepoints or crosses we could still process all populations at the same time. Furthermore, these three Selected and three Control populations were fixed for alternative alleles at candidate SNPs associated with *fiz* and showed clear differences in *fiz* expression, which was not the case for some of the populations left out of this study, notably C1 and C3 (see Results).

Before each experiment, we reared all populations on standard diet for at least two generations (relaxed selection) to limit environmental maternal effects. To obtain larvae for each experiment, we let approximately 200 adult flies from a given population to lay eggs overnight on an orange juice-agar plates supplemented with yeast. The desired number of eggs was then transferred onto the experimental media (see below) and inoculated with feces suspension (OD_600_ = 0.5) from a pool of flies from all populations to ensure homogeneity of larval microbiota (Cavigliasso, et al. 2023). The assays reported here were performed after 258 to 290 generations of experimental evolution.

### Expression of ecdysone oxidase and ecdysone-dependent genes across development

We used RT-qPCR to compare the expression of ecdysone oxidase genes and several other genes of interest between the Control (C2, C4, C6) and Selected (S1, S2, S3) populations in two independent experiments. For both experiments, the food was homogenized with a blender before adding eggs to facilitate subsequent larvae collection at different time points.

The first experiment aimed to test for consistent differences in expression throughout much of the development and into the adult stage. From each population we transferred approximately 100 eggs into each of four bottles (three bottles for larvae – one bottle per collection day; one bottle for prepupae and adults) with about 15 mL of either poor or standard diet. From the poor diet, we collected larvae at 4 and 5 days post egg laying as well as at day 7 for Controls only (at day 7, Selected larvae have already started to pupariate). From the standard diet larvae were sampled at 3, 4 and 5 days post egg-laying. From both diets we also sampled white prepupae with everted spiracles and virgin male and female adult flies (two samples of seven individuals per population, diet and stage). For each timepoint we examined another sample of larvae to determine if they were in the second or third instar (based on the shape of anterior spiracles). See Supplementary Figure S1 for the correspondence between larval stage and age.

In the second experiment we aimed to test for differences in gene expression during the first three quarters of the third larval instar. This was only done on the standard diet, which allowed for good synchronization of the developmental stage. Approximately 200 eggs were transferred to a bottle with 40 mL of standard diet. After 78-80 hours, from each bottle we sampled around 100 second instar larvae and transferred them to a new bottle containing 15 mL of fresh standard diet. 17 h later, from each bottle we collected one sample of 10 third instar larvae. At this time, we also transferred two sets of 20 3rd instar larvae in two new bottles with standard diet, and collected samples of 10 larvae 24 and 36 hours later. Three replicate samples per population and stage were collected from different sets of bottles. In this experiment, we also measured the expression of two other paralogs of *fiz* (*CG9521* and *CG12539*), two genes involved in activation/deactivation of 20E (*shade* and *Cyp18a1*) as well as two ecdysone-dependent genes (*broad* and *Eip74EF*) (see Supplementary Table S5 for primer sequences).

The samples were flash-frozen in liquid nitrogen and stored at –80°C. We extracted RNA with the RNeasy Plus mini Kit (#74134, Qiagen, Switzerland) following manufacturer’s protocol and quantified with a spectrophotometer (DS-11 FX, DeNovix, Bucher Biotec AG, Switzerland); 300 ng was used as template for the reverse transcription using the PrimeScript^TM^ RT Master mix (#RR036A, Takara, France). The qPCR was performed with a QuantStudio 6 Flex system equipped with a 384-well block and the SspAdvanced Unversal SYBR® Green Supermix (#1725272, BioRad, Switzerland) under the following conditions: 95°C for 30 sec, 40 cycles of 95°C for 15 sec and 60°C for 30 sec. The melt-curve analysis was between 65 and 95°C with a 0.5°C increment at 5 sec per step. Primer efficiency was tested on the same device. All PCR primers used in this paper were designed to avoid any SNPs (candidate or not) previously identified in these populations (Kawecki, et al. 2021). In addition to the target genes, we amplified three reference genes (*αTub84B, eEF1α2, RpL32*), chosen for their stability with diets (Ponton, et al. 2011). For the analysis of gene expression, we took the median of the technical replicates. Two to four technical replicates were done depending on the experiment and genes.

We quantified the expression of each gene (GOI) of interest relative to the (unweighted) geometric mean of the three reference genes (REF) and expressed it on log_2_ scale:

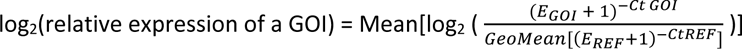

where E is the primer efficiency, *Ct* is the number of cycles until the threshold, and Mean stands for arithmetic mean.

To compare the relative expression of genes of interest (log_2_ transformed), we fitted for each gene a linear mixed model (LMM) with Type 3 F-tests and using R function *lmer* (Kuznetsova, et al. 2017). The selection regime (Control or Selected), the developmental stage and their interaction were included as fixed factors. The replicate populations nested within the selection regime in interaction with stage were included as random factors. Pairwise comparisons were performed with *emmeans* and *pairs* functions in R. P-values of pairwise comparisons were adjusted with sequential Bonferroni (Holm 1979).

### Allele-specific expression of fiz and CG9512 in F1 female larvae

To test for *cis*- or *trans*-regulatory nature of difference in *fiz* and *CG9512* expression, we used amplicon sequencing to assess the relative contribution of the alleles originating from the Selected and Control populations to mRNA pool in heterozygous larvae (Signor and Nuzhdin 2018). With *fiz* and *CG9512* being located on the X chromosome, these larvae had to be female. Because phenotypic sexing of live larvae, based on presence or absence of developing testes, is only practical in late third larval instar, we focused on this stage. We generated three types of SEL × CTL F1 crosses by crossing virgin females of populations S1, S2 and S3 with males from Control populations C2, C4 and C6, respectively. Similarly, CTL × SEL F1 crosses were obtained by making the same crosses in the reverse direction. The F1 larvae were raised on standard diet as described above; larvae of the parental Control and Selected populations were raised in parallel.

We collected samples of seven late (but not yet wandering) third instar larvae (5 days-old) judged to be female based on the apparent absence of developing testes (four samples per population or cross; four samples of male larvae were also collected for the assay described in the next subsection). The samples were flash-frozen and stored at −80°C. We extracted both RNA and DNA from these presumed-female larval samples. The samples were homogenized in 1 mL of TRIzol^TM^ Reagent (#15596018, ThermoFisher Scientific, Switzerland) solution and both DNA and RNA were extracted from each sample using the manufacturer’s recommendations. The quality of nucleic acids was assessed with a Fragment Analyzer (Agilent, Switzerland) at the Lausanne Genomic Technologies Facility of the University of Lausanne.

Sexing of larvae as female is prone to errors because it based on not seeing developing testes, which can be poorly visible (e.g., embedded in the fat body). Thus, to ensure the samples only contained female larvae, we tested for the presence of the Y chromosome by performing PCR on DNA extracted from those samples with primers specific to two Y chromosome genes, *FDY* and *Pp1-Y2* (Carvalho, et al. 2015; Dupim, et al. 2018). A further PCR on the autosomal gene *Act42A* served as a positive control for the quality of DNA. Of the 48 samples, one sample where extraction failed and eight samples in which an amplicon for at least one of these two Y-chromosome genes was detected were discarded (Supplementary Table S13, Supplementary Figure S3). This resulted in two to four replicates per cross retained for amplicon sequencing.

RNA from these retained samples was reverse-transcribed into cDNA using 250 ng as a template and the PrimeScript^TM^ RT Master mix (#RR036A, Takara, France). To provide a comparison point for the allele-specific expression in F1 larvae, we also generated 50:50 mixed parental samples by mixing equal amounts of cDNA from a Selected and a Control sample, using the same population pairings as those used to generate the crosses. The relative amounts of transcripts originating from the two populations in the 50:50 mix should thus correspond to the relative differences in expression between the populations.

To generate the amplicon, PCR was performed on the cDNA with the enzyme 2X KAPA HiFi Hotstart ReadyMix (#KK2602, Roche Diagnostic, Switzerland) and the following conditions: 95°C for 3 min, 35 cycles of 98°C for 20 sec, 60°C for 15 sec and 72°C for 15 sec and a final extension step at 72°C for 1 min. cDNA libraries were prepared according to the 16S Metagenomic Sequencing Library Preparation protocol (15044223 Rev. B, Illumina). Libraries were indexed using Nextera XT Index Kit (FC-131-1002/Illumina). They were subjected to Illumina MiSeq paired-end sequencing (MiSeq v2 PE250, Illumina, Switzerland) in one lane at the Genomic Technologies Facility of the University of Lausanne. The primers used here differed from those used for qPCR (Supplementary Table S5). They contained the overhang and were located within one exon whereas for qPCR, when possible, one primer was exon spanning. *fiz* and *CG9512* primers bonded the second exon and amplified a region containing one and three SNPs, respectively. For simplicity, for *CG9512*, we did the analysis focusing on only one of the three SNPs.

After de-multiplexing, the reads were first sorted to each gene based on the sequence using a custom script. Subsequently, the reads identified as corresponding to the allele originating from the Selected and Control population were counted. This was done based on the base identity at the coding SNPs, at which the Selected and Control populations were fixed for alternative alleles: G/A at position 791 after start codon (X:14908215) for *fiz* and C/T at position 904 (X:14904032) for *CG9512* (Kawecki, et al. 2021). With at least 23,000 reads covering each SNP in any sample we obtained precise estimates of relative abundance of the allelic transcripts.

We verified that the parental populations were indeed fixed for alternative alleles at the focal coding SNPs by sequencing genomic DNA amplicons from the female larval samples from populations C2, C4, C6, S1, S2, S3. Although some alleles in each of these genomic DNA amplicons were attributed to the allele supposedly absent from the population, the highest frequency was 1.5 %, well below the lowest non-zero frequency of 1/14 = 7.1 % possible in a sample of 7 individuals. Thus, these “wrong” alleles likely represent sequencing errors.

For each SEL × CTL, CTL × SEL and mixed parental line sample, we calculated the ratio of each allele as log(number of reads allele 1/number of reads allele 2), with allele 1 and 2 being arbitrary chosen for each gene. This ratio was then analyzed with a LMM with the cross type (SEL × CTL or CTL × SEL or mixed parental line) as a fixed factor and the pairing population nested within the cross type as a random factor. Custom contrasts using emmeans R package were then applied to test if the ratio in both F1 together deviated from 50:50, which would indicate a cis-regulatory component to their difference in expression. Furthermore, we compared the ratio of the F1 larvae with those in the mixed parental lines; if the proportion in F1 larvae deviated less from 50:50 than that of the mixed samples it would indicate a contribution of a trans-regulatory component to the expression difference.

### Expression of ecdysone oxidase genes in Selected, Control and F1 male larvae

To test whether the differential expression of *fiz*, *CG9512*, *Eo* and *CG45065* between selection regime was also observed in males, we quantified (with RT-qPCR) the expression of male Selected and Control late third instar larvae. Furthermore, we quantified the expression of these genes in larvae from F1 crosses between Selected and Control populations. Because the male larvae only carried maternal X chromosome, a difference in expression between reciprocal crosses that resembled the difference between their maternal populations would indicate that the difference in expression is mediated by (*cis*- or *trans*-acting) genetic variants located on the X chromosome.

We used samples of seven male larvae collected from the same larval cultures as the female larval samples used in amplicon sequencing (see above). RNA was extracted and cDNA generated as described above. RT-qPCR was performed with SsoAdvanced Unversal SYBR® Green Supermix (#1725272, BioRad, Switzerland) and the CFX96 Touch System (BioRad, Switzerland) under the following conditions: 95°C for 30 sec, 40 cycles of 95°C for 15 sec and 60°C for 30 sec. The melt-curve analysis was between 65 and 95°C with a 0.5°C increment at 5 sec per step. We analysed the melting curves with Bio-Rad CFX Maestro software and ensured a single product was amplified. We measured the expression of *fiz* and *CG9512* relative to three reference genes (*αTub84B, eEF1α2, RpL32,* Supplementary Table S5). The relative expression of the genes of interest (log_2_ transformed) were calculated as in 4.4. section and similar LMM were done for each gene with the population type (the selection regime (Control or Selected) and the type of F1 (CTL×SEL or SEL×CTL) as fixed factor. The replicate populations nested within the population type were included as random factor. Pairwise comparisons were performed with *emmeans* and *pairs* functions in R.

### Effect of fiz knockdown on larval performance

To knockdown *fiz* expression, we expressed a UAS-*fiz*^RNAi^ construct under the control of ubiquitously expressed Act-GAL4 driver. We crossed >100 virgin Act-GAL4/CyO, *twi*-UAS-GFP females with 20-40 males homozygous for UAS-*fiz*^RNAi^ (ID: 107089; Vienna Drosophila Resource Center, Austria). As the control, we used offspring from the cross between Act-GAL4/CyO, *twi*-UAS-GFP females and males from a line containing an empty vector at the same genomic location (UAS^−^; ID: 60100; Vienna Drosophila Resource Center, Austria). The female driver line being heterozygous, only half of the offspring would carry the driving Act-GAL4 haplotype; the other half would carry CyO, *twi*-UAS-GFP and thus express GFP. We thus sorted embryos based on the GFP phenotype (6 h after the end of oviposition to give them enough time for the GFP to become visible) and transferred groups of 200 non-GFP eggs to bottles with either standard or poor diet (N = 4-5 replicate bottles per cross and diet; for logistic reasons, the two diets were tested in separate experiments).

For each bottle of each cross and diet, we estimated the egg-to-adult survival probability (i.e., the number of eclosed adults divided by the number of eggs) and analyzed it by fitting a generalized linear mixed model (GLMM) for each diet with a binomial error distribution with the number of eclosed *versus* non-eclosed flies as the response variable, the genotype (control or *fiz*-knockdown) as fixed factor and the bottles as random factor. GLMM were fitted using the mixed function in afex R package (Singmann, et al. 2022) and significance of fixed factor was tested with Likelihood Ratio test (LRT) method. Eclosing adults of either sex were scored daily to estimate egg-to-adult developmental time (we used the inverse of the development time for the analysis). Females eclosed on each day were collected, dried at 70°C for 24h and weighed as a group with a precision balance (Mettler Toledo, MT5) to the nearest 1 μg. Growth rate was calculated as in Cavigliasso, et al. (2023) with this formula: growth rate = ln(female weight/egg weight)/larval developmental time, where egg weight was assumed to be 5 µg (Kolss, et al. 2009) and the larval developmental time was estimated as the time from egg laying to the day on which the females were collected minus 5 days to account for the time needed for embryonal development and metamorphosis. To analyze the effect of fiz-knockdown on these performances, we fitted a LMM for each diet with inverse of developmental time or female weight or female growth rate as response variable, the genotype (control *versus fiz*-knockdown) and when applicable, the sex or the developmental time, quadratic development time and their interaction with genotype as fixed factors. For all these models, the replicate bottles were used as random factors. We tested for studentized residuals and removed one outlier data point (studentized residual > 3) for weight and growth rate.

In each experiment an additional bottle with non-GFP embryos on standard diet was prepared to test the effectiveness of the knockdown. Five days later, late third instar larvae were collected in three replicates and *fiz* expression was measured with RT-qPCR (Supplementary Figure S4) as described above, using *αTub84B, eEF1α2* and *RpL32* as reference genes (Supplementary Table S5). As previously, to test the effectiveness of *fiz*-knockdown in each block, we fitted a LM and a type 2 F-test with the genotype (control *versus fiz*-knockdown) as fix factor.

## Supplementary material

Supplementary data are available at *Molecular Biology and Evolution* online.

## Supporting information

Original data

## Acknowledgements

This work was supported by the Swiss National Science Foundation (grants 31003A_162732 and 310030_184791 to TJK) and by funding from the University of Lausanne. The authors thank C. Palazzo and H. Richter for help with experiments, J. Weber and J. Marquis for advice on amplicon sequencing, M.-C. Gambetta for providing the GAL4 line and R. Arguello for comments on a previous version of the manuscript.

## Authors contributions

TJK, FC, BE, HG-A, JI and VLK conceptualized the research; FK, MS AK, BE, LS and HG-A performed the experiments; FC, HG-A and TJK curated and analyzed the data; FC and TJK visualized the results and wrote the paper with contributions and revisions from other authors.

## Data availability

Data are available as part of supplementary material.

## References

Auer TO, Shahandeh MP, Benton R. 2021. *Drosophila sechellia*: A genetic model for behavioral evolution and neuroecology. Annual Review of Genetics 55:527–554.

Baker KD, Warren JT, Thummel CS, Gilbert LI, Mangelsdorf DJ. 2000. Transcriptional activation of the *Drosophila* ecdysone receptor by insect and plant ecdysteroids. Insect Biochemistry and Molecular Biology 30:1037–1043.

Carvalho AB, Vicoso B, Russo CA, Swenor B, Clark AG. 2015. Birth of a new gene on the Y chromosome of *Drosophila melanogaster*. Proceedings of the National Academy of Sciences of the United States of America 112:12450–12455.

Cavigliasso F, Dupuis C, Savary L, Spangenberg JE, Kawecki TJ. 2020. Experimental evolution of post- ingestive nutritional compensation in response to a nutrient-poor diet. Proceedings of the Royal Society B: Biological Sciences 287:20202684.

Cavigliasso F, Savary L, Spangenberg JE, Gallart-Ayala H, Ivanisevic J, Kawecki TJ. 2023. Experimental evolution of metabolism under nutrient restriction: enhanced amino acid catabolism and a key role of branched-chain amino acids. Evolution Letters 7:273–284.

Chan YF, Marks ME, Jones FC, Villarreal G, Shapiro MD, Brady SD, Southwick AM, Absher DM, Grimwood J, Schmutz J, et al. 2010. Adaptive evolution of pelvic reduction in sticklebacks by recurrent deletion of a *Pitx1* enhancer. Science 327:302–305.

Colombani J, Bianchini L, Layalle S, Pondeville E, Dauphin-Villemant C, Antoniewski C, Carré C, Noselli S, Léopold P. 2005. Antagonistic actions of ecdysone and insulins determine final size in *Drosophila*. Science 310:667–670.

Colosimo PF, Hosemann KEB, S., Villarreal G, Dickson M, Grimwood J, Schmutz J, Myers RM, Schluter D, Kingsley DM. 2005. Widespread parallel evolution in sticklebacks by repeated fixation of ectodysplasin alleles. Science 307:1928–1933.

Cosme LV, Gloria-Soria A, Caccone A, Powell JR, Martins AJ. 2020. Evolution of kdr haplotypes in worldwide populations of *Aedes aegypti*: Independent origins of the F1534C kdr mutation. PLOS Neglected Tropical Diseases 14:e0008219.

Delanoue R, Slaidina M, Leopold P. 2010. The steroid hormone ecdysone controls systemic growth by repressing dMyc function in *Drosophila* fat cells. Develeopmental Cell 18:1012–1021.

Dreos R, Ambrosini G, Perier RC, Bucher P. 2015. The Eukaryotic Promoter Database: expansion of EPDnew and new promoter analysis tools. Nucleic Acids Research 43:D92–96.

Dupim EG, Goldstein G, Vanderlinde T, Vaz SC, Krsticevic F, Bastos A, Pinhao T, Torres M, David JR, Vilela CR, et al. 2018. An investigation of Y chromosome incorporations in 400 species of *Drosophila* and related genera. PLoS Genetics 14:e1007770.

Erkosar B, Kolly S, van der Meer JR, Kawecki TJ. 2017. Adaptation to chronic nutritional stress leads to reduced dependence on microbiota in *Drosophila melanogaster*. mBio 8.

Glaser-Schmitt A, Catalan A, Parsch J. 2013. Adaptive divergence of a transcriptional enhancer between populations of *Drosophila melanogaster*. Philosophical Transactions of the Royal Society B 368:20130024.

Glaser-Schmitt A, Parsch J. 2018. Functional characterization of adaptive variation within a cis- regulatory element influencing *Drosophila melanogaster* growth. PLoS Biology 16:e2004538.

Glaser-Schmitt A, Wittmann MJ, Ramnarine TJS, Parsch J. 2021. Sexual antagonism, temporally fluctuating selection, and variable dominance affect a regulatory polymorphism in *Drosophila melanogaster*. Molecular Biology and Evolution 38:4891–4907.

Grenier T, Consuegra J, Ferrarini MG, Akherraz H, Bai L, Dusabyinema Y, Rahioui I, Da Silva P, Gillet B, Hughes S, et al. 2023. Intestinal GCN2 controls Drosophila systemic growth in response to Lactiplantibacillus plantarum symbiotic cues encoded by r/tRNA operons. eLife 12:e76584.

Guittard E, Blais C, Maria A, Parvy JP, Pasricha S, Lumb C, Lafont R, Daborn PJ, Dauphin-Villemant C. 2011. CYP18A1, a key enzyme of *Drosophila* steroid hormone inactivation, is essential for metamorphosis. Developmental Biology 349:35–45.

Hardy CM, Burke MK, Everett LJ, Han MV, Lantz KM, Gibbs AG. 2018. Genome-Wide Analysis of Starvation-Selected Drosophila melanogaster-A Genetic Model of Obesity. Molecular Biology and Evolution 35:50–65.

Hoedjes KM, Kostic H, Flatt T, Keller L. 2023. A single nucleotide variant in the PPARgamma-homolog Eip75B affects fecundity in *Drosophila*. Molecular Biology and Evolution 40.

Hoedjes KM, van den Heuvel J, Kapun M, Keller L, Flatt T, Zwaan BJ. 2019. Distinct genomic signals of lifespan and life history evolution in response to postponed reproduction and larval diet in Drosophila. Evolution Letters 3:598–609.

Holm S. 1979. A simple sequentially rejective multiple test procedure. The Scandinavian Journal of Statistics 6:65–70.

Hubbard JK, Uy JA, Hauber ME, Hoekstra HE, Safran RJ. 2010. Vertebrate pigmentation: from underlying genes to adaptive function. Trends in Genetics 26:231–239.

Hutter S, Saminadin-Peter SS, Stephan W, Parsch J. 2008. Gene expression variation in African and European populations of *Drosophila melanogaster*. Genome Biology 9:R12.

Jakšić AM, Karner J, Nolte V, Hsu SK, Barghi N, Mallard F, Otte KA, Svečnjak L, Senti KA, Schlötterer C. 2020. Neuronal function and dopamine signaling evolve at high temperature in *Drosophila*. Molecular Biology and Evolution 37:2630–2640.

Kannangara JR, Mirth CK, Warr CG. 2021. Regulation of ecdysone production in *Drosophila* by neuropeptides and peptide hormones. Open Biology 11:200373.

Kapali GP, Callier V, Gascoigne SJL, Harrison JF, Shingleton AW. 2022. The steroid hormone ecdysone regulates growth rate in response to oxygen availability. Scientific Report 12:4730.

Kapun M, Barron MG, Staubach F, Obbard DJ, Wiberg RAW, Vieira J, Goubert C, Rota-Stabelli O, Kankare M, Bogaerts-Marquez M, et al. 2020. Genomic analysis of European *Drosophila melanogaster* populations reveals longitudinal structure, continent-wide selection, and previously unknown DNA viruses. Molecular Biology and Evolution 37:2661–2678.

Karim FD, Thummel CS. 1991. Ecdysone coordinates the timing and amounts of *E74A* and *E74B* transcription in *Drosophila*. Genes and Development 5:1067–1079.

Kawecki TJ, Erkosar B, Dupuis C, Hollis B, Stillwell RC, Kapun M. 2021. The genomic architecture of adaptation to larval malnutrition points to a trade-off with adult starvation resistance in *Drosophila*. Molecular Biology and Evolution 38:2732–2749.

Kolss M, Vijendravarma RK, Schwaller G, Kawecki TJ. 2009. Life-history consequences of adaptation to larval nutritional stress in *Drosophila*. Evolution 63:2389–2401.

Koyama T, Mirth CK. 2018. Unravelling the diversity of mechanisms through which nutrition regulates body size in insects. Current Opinion in Insect Science 25:1–8.

Krause SA, Overend G, Dow JAT, Leader DP. 2022. FlyAtlas 2 in 2022: enhancements to the *Drosophila melanogaster* expression atlas. Nucleic Acids Research 50:D1010–D1015.

Kuznetsova A, Brockhoff PB, Christensen RHB. 2017. lmerTest package: tests in linear mixed effects models. Journal of Statistical Software 82.

Lavrynenko O, Rodenfels J, Carvalho M, Dye NA, Lafont R, Eaton S, Shevchenko A. 2015. The ecdysteroidome of *Drosophila*: influence of diet and development. Development 142:3758–3768.

Lee GJ, Han G, Yun HM, Lim JJ, Noh S, Lee J, Hyun S. 2018. Steroid signaling mediates nutritional regulation of juvenile body growth via IGF-binding protein in *Drosophila*. Proceedings of the National Academy of Sciences of the United States of America 115:5992–5997.

Lin C, Koval A, Tishchenko S, Gabdulkhakov A, Tin U, Solis Gonzalo P, Katanaev Vladimir L. 2014. Double suppression of the Gα protein activity by RGS proteins. Molecular Cell 53:663–671.

Marden JH, Fescemyer HW, Schilder RJ, Doerfler WR, Vera JC, Wheat CW. 2013. Genetic variation in HIF signaling underlies quantitative variation in physiological and life-history traits within lowland butterfly populations. Evolution 67:1105–1115.

Martins NE, Faria VG, Nolte V, Schlötterer C, Teixeira L, Sucena É, Magalhães S. 2014. Host adaptation to viruses relies on few genes with different cross-resistance properties. Proceedings of the National Academy of Sciences of the United States of America 111:15597–15597.

Meylan P, Dreos R, Ambrosini G, Groux R, Bucher P. 2020. EPD in 2020: enhanced data visualization and extension to ncRNA promoters. Nucleic Acids Research 48:D65–D69.

Narasimha S, Kolly S, Sokolowski MB, Kawecki TJ, Vijendravarma RK. 2015. Prepupal building behavior in *Drosophila melanogaster* and its evolution under resource and time constraints. PLoS One 10:e0117280.

O’Brown NM, Summers BR, Jones FC, Brady SD, Kingsley DM. 2015. A recurrent regulatory change underlying altered expression and Wnt response of the stickleback armor plates gene EDA. eLife 4:e05290.

Petryk A, Warren JT, Marqués G, Jarcho MP, Gilbert LI, Kahler J, Parvy J-P, Li Y, Dauphin-Villemant C, O’Connor MB. 2003. Shade is the *Drosophila* P450 enzyme that mediates the hydroxylation of ecdysone to the steroid insect molting hormone 20-hydroxyecdysone. Proceedings of the National Academy of Sciences of the United States of America 100:13773–13778.

Ponton F, Chapuis MP, Pernice M, Sword GA, Simpson SJ. 2011. Evaluation of potential reference genes for reverse transcription-qPCR studies of physiological responses in *Drosophila melanogaster*. Journal of Insect Physiology 57:840–850.

Prentice AM. 2005. Starvation in humans: evolutionary background and contemporary implications. Mechanisms of Ageing and Development 126:976–981.

Rees H. 2013. Edcdysteroid biosynthesis and inactivation in relation to function. European Journal of Entomology 92:9–39.

Rewitz KF, Yamanaka N, O’Connor MB. 2013. Developmental checkpoints and feedback circuits time insect maturation. Current Topics in Developmental Biology 103:1–33.

Rewitz KF, Yamanaka N, O’Connor MB. 2010. Steroid hormone inactivation is required during the juvenile-adult transition in *Drosophila*. Developmental Cell 19:895–902.

Scanlan JL, Robin C, Mirth CK. 2023. Rethinking the ecdysteroid source during *Drosophila* pupal-adult development. Insect Biochemistry and Molecular Biology 152:103891.

Signor SA, Nuzhdin SV. 2018. The evolution of gene expression in cis and trans. Trends in Genetics 34:532–544.

Singmann H, Bolker B, Westfall J, Aust F, Ben-Shachar M. 2022. afex: analysis of factorial experiments. R package version 1.1–1.

Sommé-Martin G, Colardeau J, Beydon P, Blais C, Lepesant JA, Lafont R. 1990. P1 gene expression in *Drosophila* larval fat body: intuction by various ecdysteroids. Archives of Insect Biochemistry and Physiology 15:43–56.

Sommé-Martin G, Colardeau J, Lafont R. 1988. Conversion of ecdysone and 20-hydroxyecdysone into 3-dehydroecdysteroids is a major pathway in third instar Drosophila melanogaster larvae. Insect Biochemistry 18:729–734.

Sun W, Shen Y-H, Qi D-W, Xiang Z-H, Zhang Z. 2012. Molecular cloning and characterization of Ecdysone oxidase and 3-dehydroecdysone-3α-reductase involved in the ecdysone inactivation pathway of silkworm, Bombyx mori. International Journal of Biological Sciences 8:125–138.

Sun W, Shen YH, Han MJ, Cao YF, Zhang Z. 2014. An adaptive transposable element insertion in the regulatory region of the EO gene in the domesticated silkworm, Bombyx mori. Molecular Biology and Evolution 31:3302–3313.

Sun W, Wang CF, Zhang Z. 2017. Transcription factor E74A affects the ecdysone titer by regulating the expression of the *EO* gene in the silkworm, Bomby mori. Biochimica et Biophysica Acta - General Subjects 1861:551–558.

Takeuchi H, Chen JH, O’Reilly DR, Turner PC, Rees HH. 2001. Regulation of ecdysteroid signaling: cloning and characterization of ecdysone oxidase: a novel steroid oxidase from the cotton leafworm, *Spodoptera littoralis*. Journal of Biological Chemistry 276:26819–26828.

Takeuchi H, Rigden DJ, Ebrahimi B, Turner PC, Rees HH. 2005. Regulation of ecdysteroid signalling during *Drosophila* development: identification, characterization and modelling of ecdysone oxidase, an enzyme involved in control of ligand concentration. Biochemical Journal 389:637–645.

Texada MJ, Malita A, Christensen CF, Dall KB, Faergeman NJ, Nagy S, Halberg KA, Rewitz K. 2019. Autophagy-mediated cholesterol trafficking controls steroid production. Developmental Cell 48:659–671 e654.

UniProt C. 2023. UniProt: the Universal Protein Knowledgebase in 2023. Nucleic Acids Research 51:D523–D531.

Vijendravarma RK, Narasimha S, Kawecki TJ. 2012a. Chronic malnutrition favours smaller critical size for metamorphosis initiation in *Drosophila melanogaster*. Journal of Evolutionary Biology 25:288–292.

Vijendravarma RK, Narasimha S, Kawecki TJ. 2012b. Evolution of foraging behaviour in response to chronic malnutrition in *Drosophila melanogaster*. Proceedings of the Royal Society B: Biological Sciences 279:3540–3546.

Vijendravarma RK, Narasimha S, Kawecki TJ. 2013. Predatory cannibalism in *Drosophila melanogaster* larvae. Nature Communications 4:1789.

Wang CF, Zhang Z, Sun W. 2018. Ecdysone oxidase and 3-dehydroecdysone-3beta-reductase contribute to the synthesis of ecdysone during early embryonic development of the silkworm. International Journal of Biological Sciences 14:1472–1482.

Wang S-s, Wang L-l, Pu Y-x, Liu J-y, Wang M-x, Zhu J, Shen Z-y, Shen X-j, Tang S-m. 2023. *Exorista sorbillans* (Diptera: Tachinidae) parasitism shortens host larvae growth duration by regulating ecdysone and juvenile hormone titers in *Bombyx mori* (Lepidoptera: Bombycidae). Journal of Insect Science 23:6.

