## Supplementary Figures and Tables for "Cis-regulatory polymorphism at *fiz* ecdysone oxidase contributes to polygenic adaptation to malnutrition in *Drosophila*"

### Supplementary material

Fanny Cavigliasso, Mikhail Savitskiy, Alexey Koval, Berra Erkosar, Loriane Savary, Hector Gallart-Ayala, Julijana Ivanisevic, Vladimir L. Katanaev, Tadeusz J. Kawecki

**Supplementary Figures S1-S5** .......... 2

**Supplementary Tables S2-S13** .......... 5

(Supplementary Table S1 is available as a separate file)

### Supplementary Figures

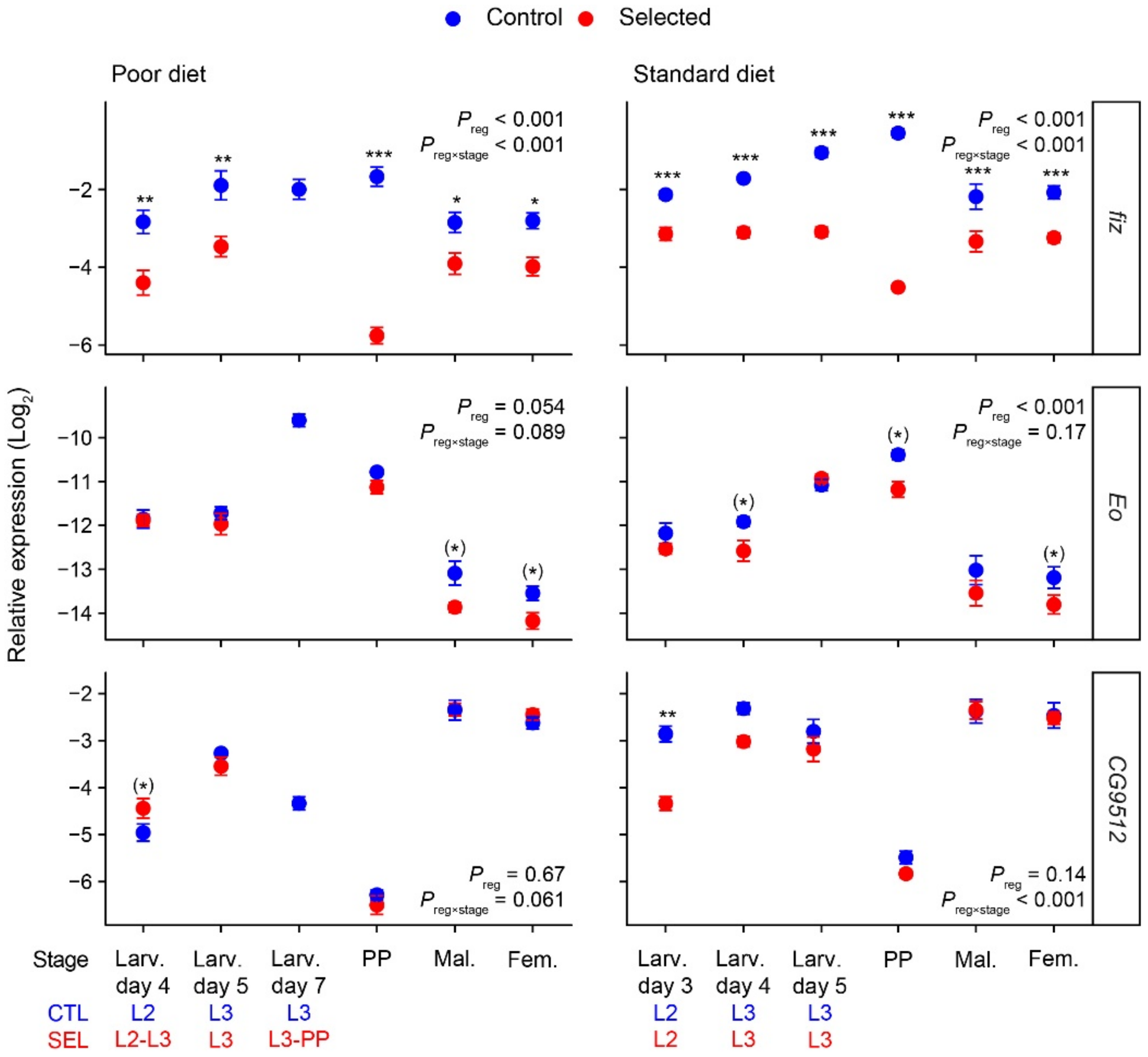


**Supplementary Figure S1.** Relative expression ($Log_2$) of *fiz, Eo* and *CG9512* according to developmental stage and diet. Larvae were collected after 3 to 7 days post egg laying. In an independent experiment, we checked the corresponding stage for Selected (SEL) and Control (CTL) populations. The correspondence is written in blue (Control) or red (Selected) below. At day 4 on poor diet, there were only L2 for CTL whereas there were ¾ of L2 and ¼ of L3 for SEL. At day 7 on poor diet, there is no data for Selected larvae because most of them pupariated by then. L2 and L3: second and third instar larvae. PP: prepupae. Fem.: Females. Mal.: Males. The expression of each target gene is quantified relative to the geometric mean of three housekeeping reference genes (*RpL32* (FBgn0002626), *αTub84B* (FBgn0003884), *eEF1α2* (FBgn0000557), Supplementary Table S5). For each gene and diet, the significance of the main effect of the selection regime ($P_{reg}$) and of the regime × stage interaction ($P_{reg×stage}$) are shown. Asterisks indicate a significant difference after p-value correction (sequential Bonferroni adjustment for stages) between Control and Selected populations at this developmental stage. Parentheses around the asterisk, it means the difference was significant before p-value correction but not after. Symbols indicate means ± SE. N = 3 populations × 2 replicates per selection regime, diet and stage. Each replicate being a pool of seven individuals (larvae, prepupae or adults). An additional model with only prepupae and adults was done for *Eo*, confirming the overall higher expression in Controls (regime on poor diet: $F_{1,6} = 12.3$, $P = 0.013$, regime on standard diet: $F_{1,6} = 8.7$, $P = 0.026$).

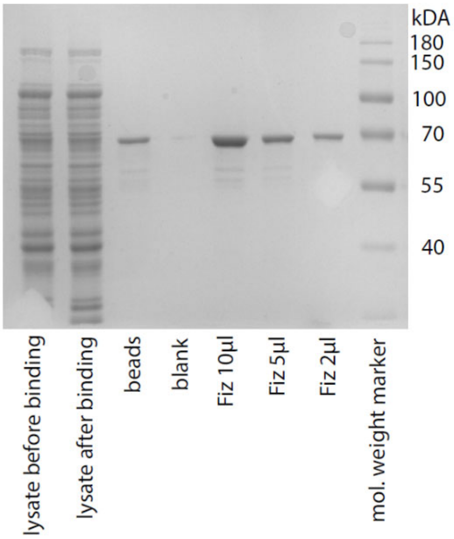


**Supplementary Figure S2.** Gel of Fiz purification.

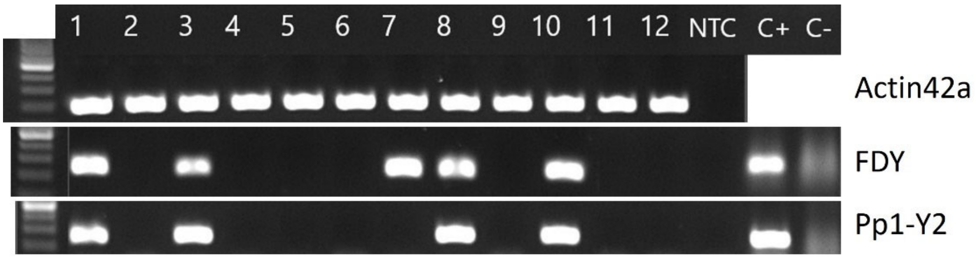


**Supplementary Figure S3.** Verification that samples for allele-specific *fiz* expression consist exclusively of female larvae (results for the first 12 samples shown). This was based on the absence of PCR amplicon for two Y chromosome genes *FDY* (FBgn0265047) and *Pp1-Y2* (FBgn0046698). C+ and C-: positive and negative controls for *FDY* and *Pp1-Y2*, corresponding to DNA pool from six adult females and one adult male or DNA pool of seven adult females, respectively. NTC: negative control without DNA. Samples with a band for *FDY* and/or *Pp1-Y2* (here samples 1, 3, 7, 8, and 10) must have contained at least one male and were excluded. A summary result is presented in Supplementary Table S13.

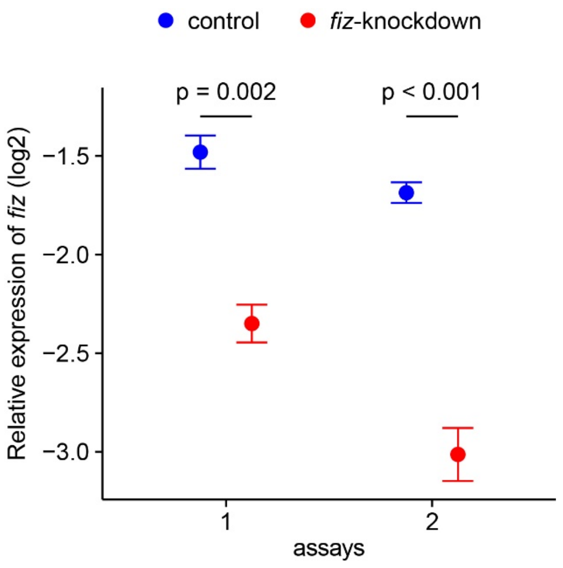


**Supplementary Figure S4.** Relative expression of *fiz* in *fiz*-knockdown and control larvae. Eggs from assay 1 was used for experiment on poor diet; eggs from assay 2 for experiment on standard diet. In both assays, the knockdown was efficient and *fiz* expression was reduced by approximately 40% (assay 1: $F_{1,4} = 46.5$, $P = 0.002$; assay 2: $F_{1,4} = 85.0$, $P < 0.001$). Symbols indicate means ± SE. N = 3 replicates of 10 larvae.

### Supplementary Tables

**Supplementary Table S1.** Allele frequencies of all SNPs found in and around *fiz* (4000 bp upstream from start codon and 2000 bp downstream from stop codon). A separate file.

**Supplementary Table S2.** Summary of significance tests from linear mixed models (LMM) for the relative amount of $H_2O_2$ in samples with purified Fiz + substrate or purified Fiz only or substrate only. Pairwise comparisons are also shown. Fiz = purified Fiz protein; E = ecdysone substrate; 20E = 20-hydroxyecdsone substrate. Results are reported in Figure 2.

| Substrate | Factor | Contrast | Estimate | Statistics | *P* |
|---|---|---|---|---|---|
| Ecdysone | Type of sample | | | $F_{2,9}$ = 1431.4 | <0.001 |
| | | (Fiz + E) – Fiz | 32512 | $t_9$ = 40.3 | <0.001 |
| | | (Fiz + E) – E | 40846 | $t_9$ = 50.6 | <0.001 |
| | | Fiz – E | 8334 | $t_9$ = 10.3 | <0.001 |
| 20E | Type of sample | | | $F_{2,4}$ = 1537 | <0.001 |
| | | (Fiz + 20E) – Fiz | 22593 | $t_4$ = 38.3 | <0.001 |
| | | (Fiz + 20E) – 20E | 21296 | $t_4$ = 51.0 | <0.001 |
| | | Fiz – 20E | -1298 | $t_4$ = -2.2 | 0.18 |

**Supplementary Table S3.** Summary of significance tests from LMM of relative expression of *fiz*, *Eo* and *CG9512* in larvae, prepupae and flies, reported in Supplementary Figure S1. The reported *P* values for contrasts are after sequential Bonferroni correction for multiple stages.

| | Poor diet | | | | Standard diet | | | |
|---|---|---|---|---|---|---|---|---|
| Gene | Factor | Stage contrast CTL – SEL | Statistics | *P* | Factor | Stage contrast CTL – SEL | Statistics | *P* |
| *fiz* | Regime | | $F_{1,6} = 90.6$ | < 0.001 | Regime | | $F_{1,4} = 340.7$ | < 0.001 |
| | Stage | | $F_{4,54} = 5.6$ | < 0.001 | Stage | | $F_{5,20} = 4.4$ | 0.007 |
| | Regime × Stage | | $F_{4,54} = 13.4$ | < 0.001 | Regime × Stage | | $F_{5,20} = 22.9$ | < 0.001 |
| | | | | | | Larvae day 3 | $t_{24} = 4.3$ | < 0.001 |
| | | Larvae day 4 | $t_{31} = 3.8$ | 0.002 | | Larvae day 4 | $t_{24} = 5.9$ | < 0.001 |
| | | Larvae day 5 | $t_{31} = 3.8$ | 0.002 | | Larvae day 5 | $t_{24} = 8.6$ | < 0.001 |
| | | Prepupae | $t_{31} = 10.0$ | < 0.001 | | Prepupae | $t_{24} = 16.8$ | < 0.001 |
| | | Males | $t_{31} = 2.6$ | 0.015 | | Males | $t_{24} = 4.9$ | < 0.001 |
| | | Females | $t_{31} = 2.9$ | 0.015 | | Females | $t_{24} = 4.9$ | < 0.001 |
| *Eo* | Regime | | $F_{1,6} = 5.7$ | 0.054 | Regime | | $F_{1,72} = 18.7$ | < 0.001 |
| | Stage | | $F_{4,54} = 143.9$ | < 0.001 | Stage | | $F_{5,72} = 72.5$ | < 0.001 |
| | Regime × Stage | | $F_{4,54} = 2.1$ | 0.089 | Regime × Stage | | $F_{5,72} = 1.6$ | 0.17 |
| | | | | | | Larvae day 3 | $t_{40} = 1.2$ | 0.48 |
| | | Larvae day 4 | $t_{29} = 0.1$ | 0.91 | | Larvae day 4 | $t_{40} = 2.2$ | 0.16 |
| | | Larvae day 5 | $t_{29} = 0.9$ | 0.80 | | Larvae day 5 | $t_{40} = 0.5$ | 0.62 |
| | | Prepupae | $t_{29} = 1.2$ | 0.75 | | Prepupae | $t_{40} = 2.6$ | 0.071 |
| | | Males | $t_{29} = 2.6$ | 0.066 | | Males | $t_{40} = 1.7$ | 0.27 |
| | | Females | $t_{29} = 2.2$ | 0.16 | | Females | $t_{40} = 2.1$ | 0.19 |
| *CG9512* | Regime | | $F_{1,6} = 0.2$ | 0.67 | Regime | | $F_{1,4} = 3.4$ | 0.14 |
| | Stage | | $F_{4,54} = 262.7$ | < 0.001 | Stage | | $F_{5,56} = 136.2$ | < 0.001 |
| | Regime × Stage | | $F_{4,54} = 2.4$ | 0.061 | Regime × Stage | | $F_{5,56} = 6.7$ | < 0.001 |
| | | | | | | Larvae day 3 | $t_{9} = 4.5$ | 0.009 |
| | | Larvae day 4 | $t_{40} = 2.1$ | 0.22 | | Larvae day 4 | $t_{9} = 2.1$ | 0.31 |
| | | Larvae day 5 | $t_{40} = 1.1$ | 1.00 | | Larvae day 5 | $t_{9} = 1.2$ | 1.00 |
| | | Prepupae | $t_{40} = 0.9$ | 1.00 | | Prepupae | $t_{9} = 1.1$ | 1.00 |
| | | Males | $t_{40} = 0.0$ | 1.00 | | Males | $t_{9} = 0.1$ | 1.00 |
| | | Females | $t_{40} = 0.7$ | 1.00 | | Females | $t_{9} = 0.2$ | 1.00 |

**Supplementary Table S4.** Summary of significance tests from LMM of relative expression of *fiz*, *Eo*, *CG9512*, *CG45065*, *CG9521*, *CG12539*, *Shade*, *Cyp18a1*, *Eip74EF* and *broad* in third instar larvae (synchronized) reported in Figure 3. The reported *P* values for contrasts are after sequential Bonferroni correction for multiple stages.

| Gene | Factor | Stage contrast CTL – SEL | Statistics | *P* | Gene | Factor | Stage contrast CTL – SEL | Statistics | *P* |
|---|---|---|---|---|---|---|---|---|---|
| | Regime | | $F_{1,6} = 55.9$ | < 0.001 | | Regime | | $F_{1,6} = 62.9$ | < 0.001 |
| | Stage | | $F_{2,48} = 11.2$ | < 0.001 | | Stage | | $F_{2,48} = 25.1$ | < 0.001 |
| *fiz* | Regime × Stage | | $F_{2,48} = 4.1$ | 0.023 | *Eo* | Regime × Stage | | $F_{2,48} = 0.6$ | 0.53 |
| | | Early L3 | $t_{12} = 5.3$ | < 0.001 | | | Early L3 | $t_{18} = 4.7$ | < 0.001 |
| | | 24h- older | $t_{12} = 5.5$ | < 0.001 | | | 24h- older | $t_{18} = 4.8$ | < 0.001 |
| | | 36h- older | $t_{12} = 6.6$ | < 0.001 | | | 36h- older | $t_{18} = 3.6$ | 0.002 |
| | Regime | | $F_{1,6} = 47.9$ | < 0.001 | | Regime | | $F_{1,6} = 1.0$ | 0.35 |
| | Stage | | $F_{2,48} = 8.8$ | < 0.001 | | Stage | | $F_{2,48} = 20.9$ | < 0.001 |
| *CG9512* | Regime × Stage | | $F_{2,48} = 22.0$ | < 0.001 | *CG45065* | Regime × Stage | | $F_{2,48} = 1.5$ | 0.23 |
| | | Early L3 | $t_{17} = 1.9$ | 0.069 | | | Early L3 | $t_{19} = 0.4$ | 0.68 |
| | | 24h- older | $t_{17} = 5.3$ | < 0.001 | | | 24h- older | $t_{19} = 1.3$ | 0.59 |
| | | 36h- older | $t_{17} = 7.0$ | < 0.001 | | | 36h- older | $t_{19} = 1.0$ | 0.68 |
| | Regime | | $F_{1,6} = 37.3$ | < 0.001 | | Regime | | $F_{1,6} = 72.3$ | < 0.001 |
| | Stage | | $F_{2,48} = 56.8$ | < 0.001 | | Stage | | $F_{2,48} = 0.1$ | 0.94 |
| *CG9521* | Regime × Stage | | $F_{2,48} = 6.2$ | 0.004 | *CG12539* | Regime × Stage | | $F_{2,48} = 0.1$ | 0.92 |
| | | Early L3 | $t_{18} = 2.3$ | 0.031 | | | Early L3 | $t_{18} = 4.6$ | < 0.001 |
| | | 24h- older | $t_{18} = 4.3$ | < 0.001 | | | 24h- older | $t_{18} = 5.1$ | < 0.001 |
| | | 36h- older | $t_{18} = 5.6$ | < 0.001 | | | 36h- older | $t_{18} = 4.9$ | < 0.001 |
| | Regime | | $F_{1,54} = 2.9$ | 0.095 | | Regime | | $F_{1,6} = 3.5$ | 0.11 |
| | Stage | | $F_{2,54} = 21.0$ | < 0.001 | | Stage | | $F_{2,48} = 87.2$ | < 0.001 |
| *Shade* | Regime × Stage | | $F_{2,54} = 0.0$ | 0.96 | *Cyp18a1* | Regime × Stage | | $F_{2,48} = 1.8$ | 0.17 |
| | | Early L3 | $t_{15} = 0.7$ | 0.82 | | | Early L3 | $t_{16} = 2.0$ | 0.18 |
| | | 24h- older | $t_{15} = 1.1$ | 0.82 | | | 24h- older | $t_{16} = 1.4$ | 0.36 |
| | | 36h- older | $t_{15} = 0.9$ | 0.82 | | | 36h- older | $t_{16} = 0.5$ | 0.60 |
| | Regime | | $F_{1,6} = 0.8$ | 0.39 | | Regime | | $F_{1,54} = 0.6$ | 0.44 |
| | Stage | | $F_{2,48} = 60.9$ | < 0.001 | | Stage | | $F_{2,54} = 279.8$ | < 0.001 |
| *Eip74EF* | Regime × Stage | | $F_{2,48} = 1.0$ | 0.39 | *broad* | Regime × Stage | | $F_{2,54} = 2.0$ | 0.15 |
| | | Early L3 | $t_{18} = 0.1$ | 1.00 | | | Early L3 | $t_{15} = 1.9$ | 0.23 |
| | | 24h- older | $t_{18} = 1.4$ | 0.50 | | | 24h- older | $t_{15} = 0.6$ | 1.00 |
| | | 36h- older | $t_{18} = 0.3$ | 1.00 | | | 36h- older | $t_{15} = 0.0$ | 1.00 |

**Supplementary Table S5.** Sequences of PCR primers used in the study. Primers used in amplicon sequencing include overhangs (in color).

| Gene | Direction | Sequence 5' -> 3' | Use | Source |
|---|---|---|---|---|
| *αTub84B* | forward | TGTCGCGTGTGAAACACTTC | RT-qPCR | Ponton et al. 2011 |
| | reverse | AGCAGGCGTTTCCAATCTG | RT-qPCR | |
| *eEF1α2* | forward | GCGTGGGTTTGTGATCAGTT | RT-qPCR | Ponton et al. 2011 |
| | reverse | GATCTTCTCCTTGCCCATCC | RT-qPCR | |
| *RpL32* | forward | ATGCTAAGCTGTCGCACAAATG | RT-qPCR | Ponton et al. 2011 |
| | reverse | GTTCGATCCGTAACCGATGT | RT-qPCR | |
| *fiz* | forward | TCTGAGTTGCCGGCACTATT | RT-qPCR | designed by authors |
| | reverse | CATTCACTCCTCCCGATCCT | RT-qPCR | |
| *Eo* | forward | GAGCTGTGCCAGGTGAAGG | RT-qPCR | www.flyrnai.org/flyprimerbank |
| | reverse | GGTCAGACCAAAGATTTCCGATT | RT-qPCR | |
| *CG9512* | forward | GCCGAAAGTGTGACCTTTGT | RT-qPCR | designed by authors |
| | reverse | ATGCCTGAAAGCAACAGGAT | RT-qPCR | |
| *CG45065* | forward | GGAGCCAACCGTCATACGTC | RT-qPCR | designed by authors |
| | reverse | TCCGAGATCTCCGTTTCATC | RT-qPCR | |
| *CG9521* | forward | ATGGCAAGCAAATCGCTGATA | RT-qPCR | www.flyrnai.org/flyprimerbank |
| | reverse | CGTCTCGAAAAGGACGTTATTGT | RT-qPCR | |
| CG12539 | forward | AGCTCAACCAGGTGGGATT | RT-qPCR | designed by authors |
| | reverse | GCTCCAGCTCCAATCACAAT | RT-qPCR | |
| *shade* | forward | CGCTTAATGCAGGGACTGTG | RT-qPCR | designed by authors |
| | reverse | GCTCTGGGGTAAACTGCTTG | RT-qPCR | |
| *Cyp18a1* | forward | GCTTCGATCCCAACAACATT | RT-qPCR | designed by authors |
| | reverse | TGTACCAGTTGCTCCTCGTG | RT-qPCR | |
| *broad* | forward | ACAACAACAGCCCCGACTT | RT-qPCR | designed by authors |
| | reverse | CGTTGCGCTTCTCCTCCTT | RT-qPCR | |
| *Eip74EF* | forward | GCTGCTCCACAATCTGCTTAG | RT-qPCR | designed by authors |
| | reverse | GCGGAAATGAACCTGTTGTG | RT-qPCR | |
| *Pp1-Y2* | forward | TGAGTCGGCTGGAATTAACC | PCR | designed by authors |
| | reverse | TGTCAGGAACGTCACATGGT | PCR | |
| *FDY* | forward | TTGCAAACTCGTGTGTGTTC | PCR | designed by authors |
| | reverse | GTTTGCCTAAGTTAAAGTATTGGATT | PCR | |
| *Act42A* | forward | CAGATGTGGATCTCGAAGCA | PCR | designed by authors |
| | reverse | TTCTGAAGGAGCGGAAGTGT | PCR | |
| *fiz* | forward | TCGTCGGCAGCGTCAGATGTGTATAAGAGACAG CTGCAGCACACGAACTTCAC | Amplicon sequencing | designed by authors |
| | reverse | GTCTCGTGGGCTCGGAGATGTGTATAAGAGACAG AGCAGCGACCATCTTTCATT | Amplicon sequencing | |
| *CG9512* | forward | TCGTCGGCAGCGTCAGATGTGTATAAGAGACAG CGTAACAATCGAGCCGAAAG | Amplicon sequencing | designed by authors |
| | reverse | GTCTCGTGGGCTCGGAGATGTGTATAAGAGACAG CGGACACGATGACCTCCTTA | Amplicon sequencing | |

**Supplementary Table S6.** Summary of significance tests from generalized linear mixed models (GLMM) for allele specific expression of *fiz* and *CG9512* in females reported in Figure 4A and 4B. F1 = average of F1 from both crosses (CTL × SEL and SEL × CTL); Mix = mix parental lines. Custom contrasts “F1” and “Mix” are compared to an allele frequency of 50%. In custom contrast “(CTL × SEL) – (SEL × CTL)”, allele frequency of CTL × SEL is compared to the one of SEL × CTL; “F1 – Mix”, allele frequency of all F1 (both direction of crosses) is compared to the allele frequency in mix parent lines.

| Gene | Custom Contrasts | Estimate | Statistics | *P* |
|---|---|---|---|---|
| *fiz* | F1 | -1.73 | $t_4 = 31.8$ | < 0.001 |
| | Mix | -2.11 | $t_6 = 26.6$ | < 0.001 |
| | (CTL × SEL) – (SEL × CTL) | -0.06 | $t_4 = 0.6$ | 0.61 |
| | F1 – Mix | 0.37 | $t_4 = 3.9$ | 0.017 |
| *CG9512* | F1 | -0.13 | $t_4 = 1.2$ | 0.30 |
| | Mix | -1.10 | $t_6 = 7.4$ | < 0.001 |
| | (CTL × SEL) – (SEL × CTL) | 0.01 | $t_4 = 0.1$ | 0.95 |
| | F1 – Mix | 0.97 | $t_4 = 5.5$ | 0.005 |

**Supplementary Table S7.** Summary of significance tests from LMM for *fiz* and *CG9512* expression in males reported in Figure 4C-D.

| Gene | Pairwise comparisons | Estimate | Statistics | *P* |
|---|---|---|---|---|
| *fiz* | Control – (CTL × SEL) | 0.12 | $t_{18} = 0.3$ | 0.99 |
| | Control – (SEL × CTL) | 2.48 | $t_{18} = 7.0$ | < 0.001 |
| | Control – Selected | 3.23 | $t_{18} = 9.1$ | < 0.001 |
| | (CTL × SEL) – (SEL × CTL) | 2.36 | $t_{18} = 6.6$ | < 0.001 |
| | (CTL × SEL) – Selected | 3.11 | $t_{18} = 8.8$ | < 0.001 |
| | (SEL × CTL) – Selected | 0.75 | $t_{18} = 2.1$ | 0.19 |
| *CG9512* | Control – (CTL × SEL) | 0.19 | $t_{18} = 1.3$ | 0.60 |
| | Control – (SEL × CTL) | 1.13 | $t_{18} = 7.3$ | < 0.001 |
| | Control – Selected | 1.36 | $t_{18} = 8.8$ | < 0.001 |
| | (CTL × SEL) – (SEL × CTL) | 0.93 | $t_{18} = 6.1$ | <0.001 |
| | (CTL × SEL) – Selected | 1.17 | $t_{18} = 7.6$ | <0.001 |
| | (SEL × CTL) – Selected | 0.24 | $t_{18} = 1.5$ | 0.44 |

**Supplementary Table S8.** Summary of significance tests from LMM for inverse of developmental time. Results are reported in Figure 5B. Genotype: control *versus fiz* knockdown.

| Diet | Factor | Contrasts control – *fiz* knockdown | Statistics | *P* |
|---|---|---|---|---|
| Poor | Genotype | | $F_{1,6} = 18.9$ | 0.005 |
| | Sex | | $F_{1,6} = 1.1$ | 0.33 |
| | Genotype × Sex | | $F_{1,6} = 4.5$ | 0.077 |
| | | in Females | $t_9 = 4.8$ | < 0.001 |
| | | in Males | $t_9 = 2.9$ | 0.019 |
| Standard | Genotype | | $F_{1,8} = 0.3$ | 0.59 |
| | Sex | | $F_{1,8} = 96.0$ | < 0.001 |
| | Genotype × Sex | | $F_{1,8} = 0.1$ | 0.76 |
| | | in Females | $t_8 = 0.5$ | 0.61 |
| | | in Males | $t_8 = 0.6$ | 0.56 |

**Supplementary Table S9.** Summary of significance tests from LMM for female weight. Pairwise comparisons were done from the model with developmental time as a factor and without quadratic factors (see methods). Results are reported in Figure 5C. Genotype: control *versus fiz* knockdown; Dev time: developmental time. The reported *P* values for contrasts are after sequential Bonferroni correction for multiple days.

| Diet | Factor | Contrasts control – fiz knockdown | Statistics | *P* |
|---|---|---|---|---|
| Poor | Genotype | | $F_{1,10} = 25.1$ | < 0.001 |
| | Dev time | | $F_{1,35} = 10.2$ | 0.003 |
| | (Dev time)$^2$ | | $F_{1,35} = 38.5$ | < 0.001 |
| | Genotype × Dev time | | $F_{1,35} = 0.7$ | 0.42 |
| | Genotype × (Dev time)$^2$ | | $F_{1,35} = 2.8$ | 0.10 |
| | | Day 15 | $t_{32} = 4.1$ | 0.002 |
| | | Day 16 | $t_{25} = 5.3$ | < 0.001 |
| | | Day 17 | $t_{27} = 4.1$ | 0.002 |
| | | Day 18 | $t_{25} = 4.1$ | 0.002 |
| | | Day 19 | $t_{25} = 3.4$ | 0.005 |
| | | Day 20 | $t_{27} = 3.2$ | 0.005 |
| Standard | Genotype | | $F_{1,20} = 13.0$ | 0.001 |
| | Dev time | | $F_{1,14} = 39.3$ | < 0.001 |
| | (Dev time)$^2$ | | $F_{1,14} = 0.9$ | 0.35 |
| | Genotype × Dev time | | $F_{1,14} = 4.1$ | 0.062 |
| | Genotype × (Dev time)$^2$ | | $F_{1,14} = 0.2$ | 0.70 |
| | | Day 12 | $t_{21} = -4.1$ | 0.002 |
| | | Day 13 | $t_{21} = -3.7$ | 0.003 |
| | | Day 14 | $t_{21} = -1.9$ | 0.070 |

**Supplementary Table S10.** Summary of significance tests from LMM for female growth rate. Pairwise comparisons were done from the model with developmental time as a factor and without quadratic factors (see methods). Results are reported in Figure 5D. Genotype: control *versus fiz* knockdown; Dev time: developmental time. The reported *P* values for contrasts are after sequential Bonferroni correction for multiple days.

| Diet | Factor | Contrasts control – fiz knockdown | Statistics | *P* |
|---|---|---|---|---|
| Poor | Genotype | | $F_{1,12} = 27.4$ | < 0.001 |
| | Dev time | | $F_{1,36} = 3318.2$ | < 0.001 |
| | (Dev time)$^2$ | | $F_{1,35} = 192.6$ | < 0.001 |
| | Genotype × Dev time | | $F_{1,36} = 2.1$ | 0.15 |
| | Genotype × (Dev time)$^2$ | | $F_{1,35} = 4.0$ | 0.052 |
| | | Day 15 | $t_{31} = 5.0$ | < 0.001 |
| | | Day 16 | $t_{24} = 6.9$ | < 0.001 |
| | | Day 17 | $t_{27} = 5.1$ | < 0.001 |
| | | Day 18 | $t_{24} = 4.8$ | < 0.001 |
| | | Day 19 | $t_{24} = 3.6$ | 0.003 |
| | | Day 20 | $t_{27} = 3.0$ | 0.006 |
| Standard | Genotype | | $F_{1,20} = 15.6$ | < 0.001 |
| | Dev time | | $F_{1,14} = 4472.9$ | < 0.001 |
| | (Dev time)$^2$ | | $F_{1,14} = 38.0$ | < 0.001 |
| | Genotype × Dev time | | $F_{1,14} = 9.6$ | 0.008 |
| | Genotype × (Dev time)$^2$ | | $F_{1,14} = 0.0$ | 0.85 |
| | | Day 12 | $t_{22} = 5.3$ | < 0.001 |
| | | Day 13 | $t_{21} = 4.1$ | 0.001 |
| | | Day 14 | $t_{21} = 1.8$ | 0.091 |

**Supplementary Table S11.** Comparison of the effect of *fiz*-knockdown on performances when raised on poor *versus* standard diet. The absolute estimates and standard errors are reported for each trait. *P*: p-value from t-test comparing the absolute estimate on poor *versus* standard diet.

| trait | sex | Poor diet | | Standard diet | | *P* |
|---|---|---|---|---|---|---|
| | | Absolute estimate | Standard error | Absolute estimate | Standard error | |
| Survival | both | 7.76e-01 | 1.07e-01 | 5.98e-01 | 1.10e-01 | 0.156 |
| Developmental time | female | 3.41e-03 | 7.05e-04 | 7.79e-04 | 1.48e-03 | 0.002 |
| | male | 2.02e-03 | 7.05e-04 | 8.93e-04 | 1.48e-03 | 0.002 |
| Weight | female | 3.73e-02 | 7.44e-03 | 2.81e-02 | 7.77e-03 | 0.011 |
| Growth rate | female | 1.32e-02 | 2.53e-03 | 1.02e-02 | 2.59e-03 | 0.004 |

**Supplementary Table S12**. Raw data on abundance of FAD (peak area; arbitrary units) in samples containing purified Fiz with a substrate (ecdysone or 20E) or only the substrate. n.d. not detected. The presence of FAD in samples containing Fiz suggests that Fiz, predicted to be a flavoprotein, is properly folded.

| Samples | Presence of Fiz | Substrate | FAD (peak area) |
|---|---|---|---|
| Fiz_1 | Yes | Ecdysone | 65905 |
| Fiz_2 | Yes | Ecdysone | 99808 |
| Fiz_3 | Yes | Ecdysone | 73717 |
| Fiz_4 | Yes | Ecdysone | 77037 |
| Fiz_5 | Yes | 20E | 97036 |
| Fiz_6 | Yes | 20E | 103759 |
| Fiz_7 | Yes | 20E | 97977 |
| Fiz_8 | Yes | 20E | 93297 |
| N_1 | No | Ecdysone | 518 |
| N_2 | No | Ecdysone | 698 |
| N_3 | No | Ecdysone | n.d. |
| N_4 | No | Ecdysone | 239 |
| N_5 | No | 20E | n.d. |
| N_6 | No | 20E | 674 |
| N_7 | No | 20E | 918 |
| N_8 | No | 20E | 693 |

**Supplementary Table S13.** Summary of the verification that samples for allele-specific *fiz* expression consists exclusively of female larvae. Verification is based on PCR targetting two genes located on the Y-chromosome: *FDY* (FBgn0265047) and *Pp1-Y2* (FBgn0046698). A “yes” or “no” indicate a presence or absence, respectively, of a band for the corresponding gene. See Supplementary Figure S4 for more details and as an example for the first 12 samples. A gray background indicates a discarded sample (i.e. with at least one male larva). The last column indicates samples of Selected and Control populations used to generate the 50:50 mixed samples of cDNA reported in Figure 4A and B.

| Sample | Population | Type | *FDY* | *Pp1-Y2* | Decision | cDNA_mixed |
|---|---|---|---|---|---|---|
| 1 | C2 | Control | Yes | Yes | Discard | |
| 2 | C4 | Control | No | No | Keep | Mix_B1 |
| 3 | C6 | Control | Yes | Yes | Discard | |
| 4 | S1 | Selected | No | No | Keep | |
| 5 | S2 | Selected | No | No | Keep | Mix_B1 |
| 6 | S3 | Selected | No | No | Keep | |
| 7 | C2×S1 | CTL×SEL | Yes | No | Discard | |
| 8 | C4×S2 | CTL×SEL | Yes | Yes | Discard | |
| 9 | C6×S3 | CTL×SEL | No | No | Keep | |
| 10 | S1×C2 | SEL×CTL | Yes | Yes | Discard | |
| 11 | S2×C4 | SEL×CTL | No | No | Keep | |
| 12 | S3×C6 | SEL×CTL | No | No | Keep | |
| 13 | C2 | Control | No | No | Keep | Mix_A1 |
| 14 | C4 | Control | No | No | Keep | Mix_B2 |
| 15 | C6 | Control | No | No | Keep | Mix_C1 |
| 16 | S1 | Selected | No | No | Keep | Mix_A1 |
| 17 | S2 | Selected | No | No | Keep | Mix_B2 |
| 18 | S3 | Selected | No | No | Keep | Mix_C1 |
| 19 | C2×S1 | CTL×SEL | No | No | Keep | |
| 20 | C4×S2 | CTL×SEL | No | No | Keep | |
| 21 | C6×S3 | CTL×SEL | No | No | Keep | |
| 22 | S1×C2 | SEL×CTL | No | No | Keep | |
| 23 | S2×C4 | SEL×CTL | No | No | Keep | |
| 24 | S3×C6 | SEL×CTL | No | No | Keep | |
| 25 | C2 | Control | No | No | Keep | Mix_A2 |
| 26 | C4 | Control | Yes | Yes | Discard | |
| 27 | C6 | Control | No | No | Keep | Mix_C2 |
| 28 | S1 | Selected | No | No | Keep | Mix_A2 |
| 29 | S2 | Selected | No | No | Keep | |
| 30 | S3 | Selected | No | No | Keep | Mix_C2 |
| 31 | C2×S1 | CTL×SEL | No | No | Keep | |
| 32 | C4×S2 | CTL×SEL | No | No | Keep | |
| 33 | C6×S3 | CTL×SEL | No | No | Keep | |
| 34 | S1×C2 | SEL×CTL | No | No | Keep | |
| 35 | S2×C4 | SEL×CTL | No | No | Keep | |
| 36 | S3×C6 | SEL×CTL | No | No | Keep | |
| 37 | C2 | Control | Yes | Yes | Discard | |
| 38 | C4 | Control | Yes | Yes | Discard | |
| 39 | C6 | Control | No | No | Keep | Mix_C3 |
| 40 | S1 | Selected | No | No | Keep | |
| 41 | S2 | Selected | No | No | Keep | |
| 42 | S3 | Selected | No | No | Keep | Mix_C3 |
| 43 | C4×S2 | CTL×SEL | No | No | Keep | |
| 44 | C6×S3 | CTL×SEL | No | No | Keep | |
| 45 | S1×C2 | SEL×CTL | No | No | Keep | |
| 46 | S2×C4 | SEL×CTL | No | No | Keep | |
| 47 | S3×C6 | SEL×CTL | No | No | Keep | |
