## Supplementary material for "Cis-regulatory polymorphism at *fiz* ecdysone oxidase contributes to polygenic adaptation to malnutrition in *Drosophila*": Original data: Data_ReadMe.txt

This respository contains the data for the paper "Cis-regulatory polymorphism at fiz ecdysone oxidase contributes to polygenic adaptation to malnutrition in Drosophila"

It contains the following files

Data1_Fiz_activity: 
Activity of the Fiz enzyme in vitro
Results in Figure 2

Data2_Gene_expression_throughout_development: 
Expression of fiz, Eo and CG9512 in Control and Selected populations at different stages on both diets
RT-qPCR
Results in Supplementary Figure S1

Data3_Expression_synchronized_L3: 
Expression of ecdysone oxidase genes + ecdysone-dependent genes 
in third instar larvae from Control and Selected populations, 
raised on standard diet and synchronized at L2-L3 transition
RT-qPCR
Results in Figure 3

Data4_Allele_specific_expression_female_larvae:
Allele-specific read counts in amplicons of fiz and CG9512 in heterozygous female larvae
Results in Figure 4A,B

Data5_Expresssion_in_male_larvae_qPCR:
Expression of fiz and CG9512 in male larvae from crosses between regimes (CTL x SEL; SEL x CTL; SEL; CTL)
RT-qPCR
Results in Figure 4C,D

Data6_fiz_RNAi_expression:
Expression of fiz in RNAi knockdown larvae and corresponding control larvae
RT-qPCR
Results in Supplementary Figure S5

Data7_fiz_RNAi__performance: 
Egg-to-adult survival, developmental time and female weight data of fiz knockdown (RNAi) and control flies
Results in Figure 5
